# Multiscale Analysis of Pangenome Enables Improved Representation of Genomic Diversity For Repetitive And Clinically Relevant Genes

**DOI:** 10.1101/2022.08.05.502980

**Authors:** Chen-Shan Chin, Sairam Behera, Asif Khalak, Fritz J Sedlazeck, Justin Wagner, Justin M. Zook

**Author notes:** Authors are listed in alphabetical order except the corresponding author.

## Abstract

The advancements in sequencing technologies and assembly methods enable the regular production of high-quality genome assemblies characterizing complex regions. However, challenges remain in efficiently interpreting variations at various scales, from smaller tandem repeats to megabase re-arrangements, across many human genomes. We present a pangenome research toolkit enabling analyses of complex pangenome variations at multiple scales. A graph decomposition method is developed for interpreting such variations. Surveying a set of 395 challenging and medically important genes in pangenome provides quantitative insights into repetitiveness and diversity that could impact the accuracy of variant calls. We apply the graph decomposition methods to the Y-chromosome gene, DAZ1/DAZ2/DAZ3/DAZ4, of which structural variants have been linked to male infertility, and X-chromosome genes OPN1LW and OPN1MW linked to eye disorders, highlighting the power of PGR-TK and pangenomics to resolve complex variation in regions of the genome that were previously too complex to analyze across many haplotypes.

## Introduction

Studying genomes, the fundamental information contained in all living beings, is the foundation for understanding the biology and evolution of all organisms, as well as the genetic diseases of humans. Despite the millions of human genomes that have been sequenced since the onset of the Human Genome Project^1,2^, and the dramatic reduction in the cost of short-read (~150bp) DNA sequencing, there is still fundamental information yet to be revealed in genomics^3^. While it is important to recognize successes to date, including small variant surveys, genome-wide association studies^4–7^, and the development of routine lab tests for genetic-based precision medicine ^8–11^, there remain fundamental biological questions that involve structures at longer length scales that can only be captured using long-range information accessible by long-read technologies and diploid phased assemblies^12–14^.

With the possibility of resolving variants at multiple scales, small and large, researchers now can fully characterize previously inaccessible regions by focusing on SNPs and small indels alone^15,16^. Examples of such previously inaccessible regions include centromere, telomeres, and complex repeat regions. Recent results with a pangenome-scale de novo human assemblies and the CHM13 telomere to telomere assembly have already shown the potential for revealing biological insights ^3,17–20^, which are the foundation for understanding complex genetic diseases.

A concept that becomes powerful in such analyses is that of the pangenome -- that is, a characterization of both the genetic structure and the genetic variation across diverse individuals of a species. However, such complexity and diversity generate interpretive challenges that require more advanced tools. A graph representing many genome assemblies at once provides a way to visualize and analyze complicated structural variations among different haplotypes^21–28^. Previously, different approaches to generate graphs representing genome structures have been proposed, e.g., cactus graphs^25^, stringomics graph with “stringlet”^21^, variant graph^27^, minigraph with reference based alignment^24^, graphs derived from all pairwise alignments^22^, or k-mer and de Bruijn graph based approaches^26,29^. These tools provide more accessible pictures for researchers to understand repeats and rearrangements than using computational intensive and visually complicated multiple sequence alignments^30–32^. In the meantime, while a graph is an elegant data structure for gathering information from pan-genomic assemblies, there remains a gap in projecting the underlying linear sequences onto a graph at various scales to reveal and compare features of many different haplotypes^23^.

To address this gap, we present a generalized graph framework as a software package, PanGenome Research Tool Kit (PGR-TK, https://github.com/Sema4-Research/pgr-tk), that is scalable to rapidly represent multiple samples at varying resolution levels. PGR-TK applies the computation techniques and data structures initially developed for fast genome assemblers^33–35^ to pan-genomics analysis tasks. Instead of building a whole genome graph at once, which can be computationally expensive. PGR-TK provides tools for building an indexed sequence database, fetching and querying sequences of interest (e.g. genes or regions with large scale structural variations) from the database to create pan-genomics graphs accordingly. It uses minimizer anchors to generate pangenome graphs at different scales without more computational intensive sequence-to-sequence alignment or explicitly calling variants with respect to a reference. As it does not use a reference for alignment, the generated graph is not subject to reference bias. We also developed an algorithm to decompose tangled pangenome graphs to more manageable units (principal bundles). With such decomposition, we can easily project the linear genomics sequence onto the principal bundles. It can provide more straightforward visualization to generate insight by revealing the contrast of the repeat and rearrangement variations among the haplotypes. Such pangenome-level graph decomposition provides utilities similar to the A-de Bruijn graph approach for identifying repeats and conserved segmental duplications^36–38^, but for the whole human pangenome collection at once. Finally, we provide PGR-TK as a software library so a researcher can do interactive analysis utilizing Jupyter Lab and related data science tools for complicated analysis tasks and build command line tools upon it.

We demonstrate the ability of PGR-TK to visualize complex variants in repetitive genes, including a gene within nested palindromic and tandem repeats (AMY1A), the Genome in A Bottle (GIAB) MHC benchmark (HLA Class II locus)^39^, the GIAB challenging medically relevant gene list^40^, and chrX and chrY ampliconic genes^41^. GIAB has excluded many of these genes from its benchmarks because they are challenging to represent in VCF and challenging to compare differing representations^42^. To understand how PGR-TK can help with the challenge of variant calling and comparison in these genes, we used the Human Pangenome Reference Consortium^19,43^ year one 47 human genome assemblies (94 diverse haplotypes). We generated pangenome graphs and developed two metrics (see **Results**) to evaluate the polymorphism in and the repetitiveness in the pangenome to identify the most complex medically relevant genes across the HPRC individuals. With the ability to survey a large set of genes swiftly with PGR-TK, we hope to understand how to better provide a broader benchmark set for challenging genes utilizing HPRC assemblies in the future. As GIAB is extending the benchmark beyond autosomal genes, we examine OPN1LW and OPN1MW on chromosome X and DAZ1/2/3/4 on chromosome Y in detail to understand how the limit due to complicated large scale genome rearrangement impacts the current methodology of generating variant call benchmarks. Our initial analysis of the GIAB Clinical And Medically Important Genes(CMRG) with a pangenome graph approach will help the research community to adapt the pangenome resource for clinical and medical genetic applications. Tools for visualizing and analyzing complicated re-arrangement loci such as PGR-TK will be essential for better variant calling and understanding the related mechanism for the community.

## Results

### Pangenome Research Toolkit

The PGR-TK has several different components to facilitate rapid pangenome analysis. The general scope and design of the PGR-TK is illustrated in **Figure 1a**. PGR-TK implements the most computationally intensive algorithms with the Rust programming language and exposes an application interface in the Python programming language. This enables scripting and interactive analysis, e.g., with Jupyter Notebook ^44^, and utilizes a compiled language for computation performance.

**Figure 1.**
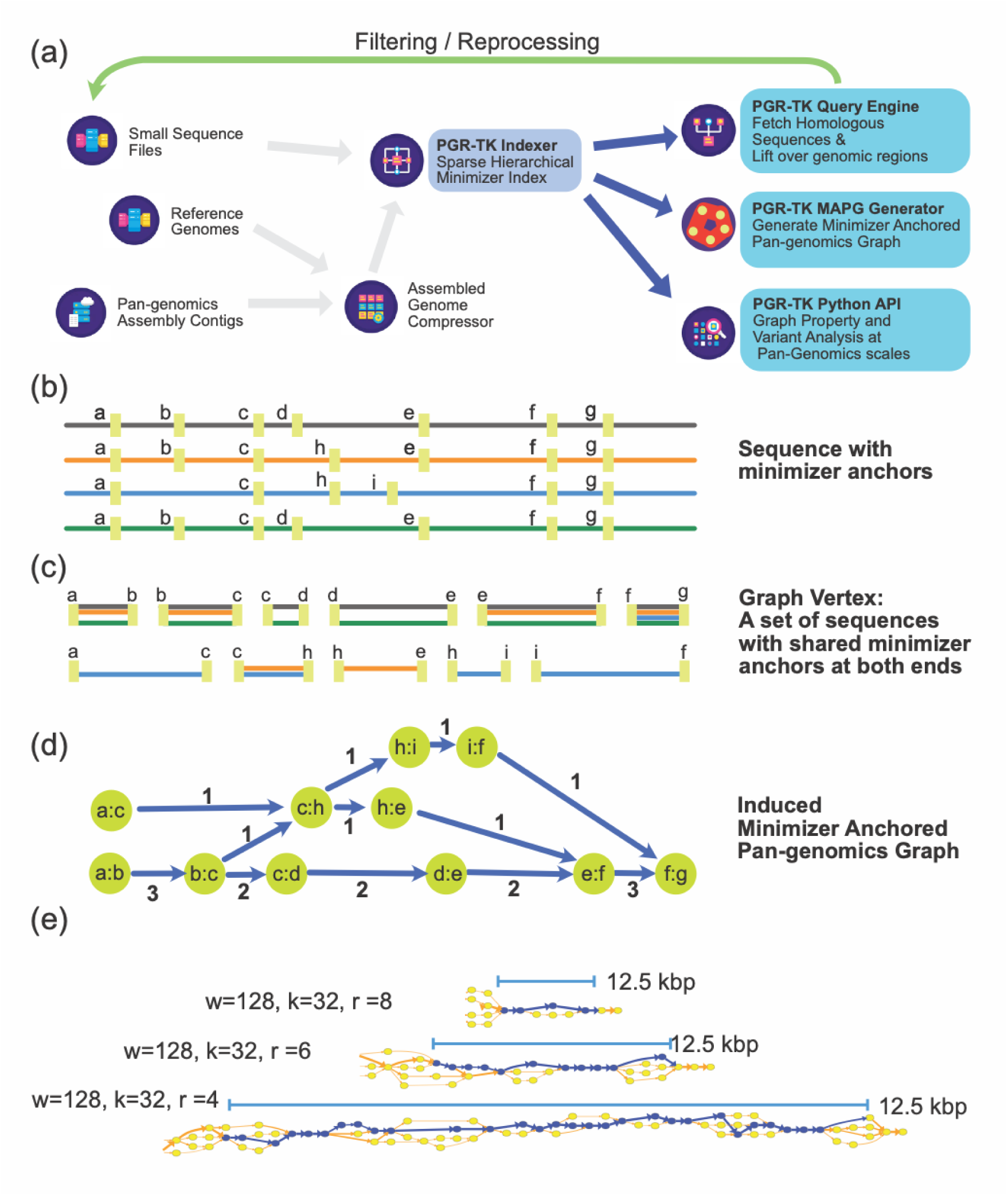
(a) Overall architecture and design scope of the PGR-TK library (b) Each sequence in the database is scanned, and the location of the minimizers are recorded to construct the SHIMMER database and minimizer anchored pangenome graph. (c) Each vertex in the MAP graph represents a collection of sequence fragments sharing the two ending SHIMMERs in the database. (d) MAP graph is constructed by merging all paths from all sequences into a graph. (d) KCNE1 MAP graph examples for 3 different scales

PGR-TK incoporated the Assembly Genome Compressor ^45^ to store pangenome assembly contigs. A command line binary is provided to create the **s**parse **hi**erarchical **m**ini**m**iz**er** (SHIMMER) index. For the HPRC year-one data release (94 fully assembled haplotypes from 47 samples), it takes 18 minutes to create the index file on an AWS c5.12xlarge instance, with the default parameters.

Once the index is built, it can be loaded into memory within minutes. As shown in **Figure 1a**, there are three main functional modules utilizing the index: (1) fetching homologous regions and sequences of the pangenome database given a query sequence, (2) creating Minimizer Anchored Pangenome (MAP) Graph, and (3) auxiliary functions for interactive analysis and visualization on the generated graph and the underlying sequences.

The source code and library can be downloaded from https://github.com/Sema4-Research/pgr-tk.

### Sparse Hierarchical Minimizer Index

Sparse Hierarchical Minimizer (SHIMMER) is a data structure extending the minimizer for more efficient indexing over larger regions. Additional minimizer reduction steps to generate sparse minimizers are applied to the minimizer sequences instead of the original base-pair sequences in a hierarchical way^33^. Such sparse minimizers can serve as natural “anchors” on genomics sequences without an explicit reference coordinate system. We utilize the SHIMMERs for quick sequence queries as initially proposed by Roberts, M. et al.^46^. PGR-TK creates an index of all neighboring pairs of SHIMMERs to the segments of sequences within the pairs.

When building a sequence index, each contig sequence is scanned and the anchoring SHIMMERs are identified. For example, **Figure 1b** shows the SHIMMERs identified on each sequence. Then, the pairs of the neighboring SHIMMERs are used for indexing the corresponding sequence segments within the paired SHIMMERs. After that, we build a look-up table of all pairs of SHIMMERs to all segments with the same pair at both ends (**Figure 1c**). For the query, we compute the neighboring SHIMMER pairs from a query sequence and search the database for all segments indexed by the same pairs. Finally, we can fetch all target segments stored in the database to get all related sequence information for further analysis. PGR-TK provides functions to refine the raw query results and filter out spurious alignments likely caused by repeats outside the region of interest. With the set of sequences homologous to the query sequence, we can quickly perform downstream analysis work, e.g., variant discovery by aligning the sequences to each other. Furthermore, we can generate a local pan-genomics (MAP) graph for comparing the sequences in the pangenome dataset at various scales by adjusting parameters to fit different analysis tasks.

### Minimizer Anchored Pangenome (MAP) Graph

The SHIMMER anchors provide a straightforward way to construct a pan-genomics graph without explicit sequence alignments or variant calling that are used in the previous pangenome graph approaches^22,24,47^. As a result, the topology of the generated pangenome graphs directly reflects the underlying genome structures in the minimizer space. It could be beneficial to avoid ambiguous cases aligning two sequences per base to each other. For example, when alignment is computed between two sequences with different numbers of tandem repeats, it may not be apparent where the additional insertions should be placed in a repeat array inside a reference sequence. Also, when constructing genomics graphs using alignment methods, the generated graph may depend on the reference used and the alignment order between the pairs of the sequences.

PGR-TK can take a set of homologous sequences to generate the “Minimizer Anchored Pangenome(MAP) graph.” The vertices in a MAP graph are labeled with the neighboring SHIMMER pairs representing a set of sequence segments in the database (**Figure 1c, 1d, and Methods**). The edges in the MAP graph are induced when at least one sequence connects the two fragments. Thus, each sequence naturally corresponds to a path in the graph, and the vertices in the path also contain the segments of other sequences in the database that share the same SHIMMER pair label. Please see the **Methods** for a precise mathematical definition of a MAP graph. The deployment of minimizer- or minhash-based approaches successfully in sequence comparison^33,34,48^ indicates that sequence segments with the same minimizer labels are also likely to be highly homologous. If necessary, we can confirm the homology or identify the variations by explicit sequence alignment of the segment inside a MAP graph vertex. The computation intensive base-to-base alignment is not required for building the MAP graph.

As the distance between minimizers can be adjusted, we can tune the parameters such that each vertex may represent segments of different sizes. This is a useful property such that we can generate a pangenome graph with different levels of details to study genomics features of different length scales. For example, the tandem repeats within the human genome may vary from a couple of hundreds of bases to 5 to 20 kilo bases. A tunable pangenome graph is useful to reveal the structures of variations within the population and to extract the related features of a specific scale.

There are three main parameters allowing us to study pan-genomics structure in different scales: (1) minimizer window size: “w”, (2) minimizer size: “k” and (3) hierarchical reduction factor: “r”. There is one more auxiliary parameter, “min_span”, that controls the minimum spanning distance between two minimizers used to construct the SHIMMER index and MAP graph (see Method).

Generally, with a given a specific minimizer size “k”, the window size “w” and the reduction factor “r” parameters, it controls the minimizer density used for the anchors. With a larger “w” and “r”, each MAP graph vertex represents bigger segment sizes. The generated graph may only capture larger differences between the different genomic sequences as the sequences are only sampled sparsely (e.g. structural variations). In such cases, the detailed differences between genomes, e.g., SNPs, are hidden inside each vertex. In contrast, if “w” and “r” are small, the generated MAP graphs contain more information for variations between sequences. **Figure 1e** shows an example of the MAP graph generated from a 12.5kbp of KCNE1 in three different scales. The same regions are represented by three MAP graphs of different scales. The sequence from the 12.5 kbp region of the hg19 KCNE1 sequences are represented with fewer nodes with r=8 than those with r=4. The r=4 graph contains more detailed information. (**Supplementary Figure 1** shows full KCNE1 MAP graphs for r from 1 to 12.) In an extreme case, when w=1, r=1, min_span=0, the generated graph is isomorphic to the de Bruijn graph^49^ with vertex symbol length size k.

### Visualizing and Analyzing Large-scale Pangenome Repeat Structures with MAP graphs

A pangenome graph can serve as a cornerstone for analyzing repeat structure variation in population^17,19^. It is usually hard to compare multiple sequences with complicated repeat structure by examining pairwise sequence alignments directly. The traditional visualization technique “dot-plot^50^” allows us to perceive the complexity of the repeats but it does not provide insights into the repeat structures as linear representations across each individual sequence directly. Furthermore, only two sequences can be compared with a dot-plot.

As an example, we use PGR-TK to investigate the repeat structure of the AMY1A (Alpha-amylase 1, an enzyme for the first step of catalyzing starch and glycogen in saliva) locus. We pick AMY1A as it has various numbers of copies caused by larger scale structure variation related to the repeat surrounding the gene. The dot-plots from randomly picking 36 sequences of a 400 kb region around AMY1A in the first year HPRC assemblies to the hg19 AMY1A reference sequence are shown in **Supplementary Figure 2a**. Visual inspections of the dot plots show there are various numbers of copies of forward repeats and inverted repeats forming palindrome sequences at the scales of 100kb, and from zero up to 5 palindrome units. Still only pairwise comparisons are enabled with dotplots and thus we lack a comprehensive assessment.

For comparison, we generate the AMY1A MAP graphs at two different scales (**Figure 2a**) from the HPRC year one assemblies (47 samples). These can be generated with PRG-TK in less than three minutes from indexed sequence data.

**Figure 2.**
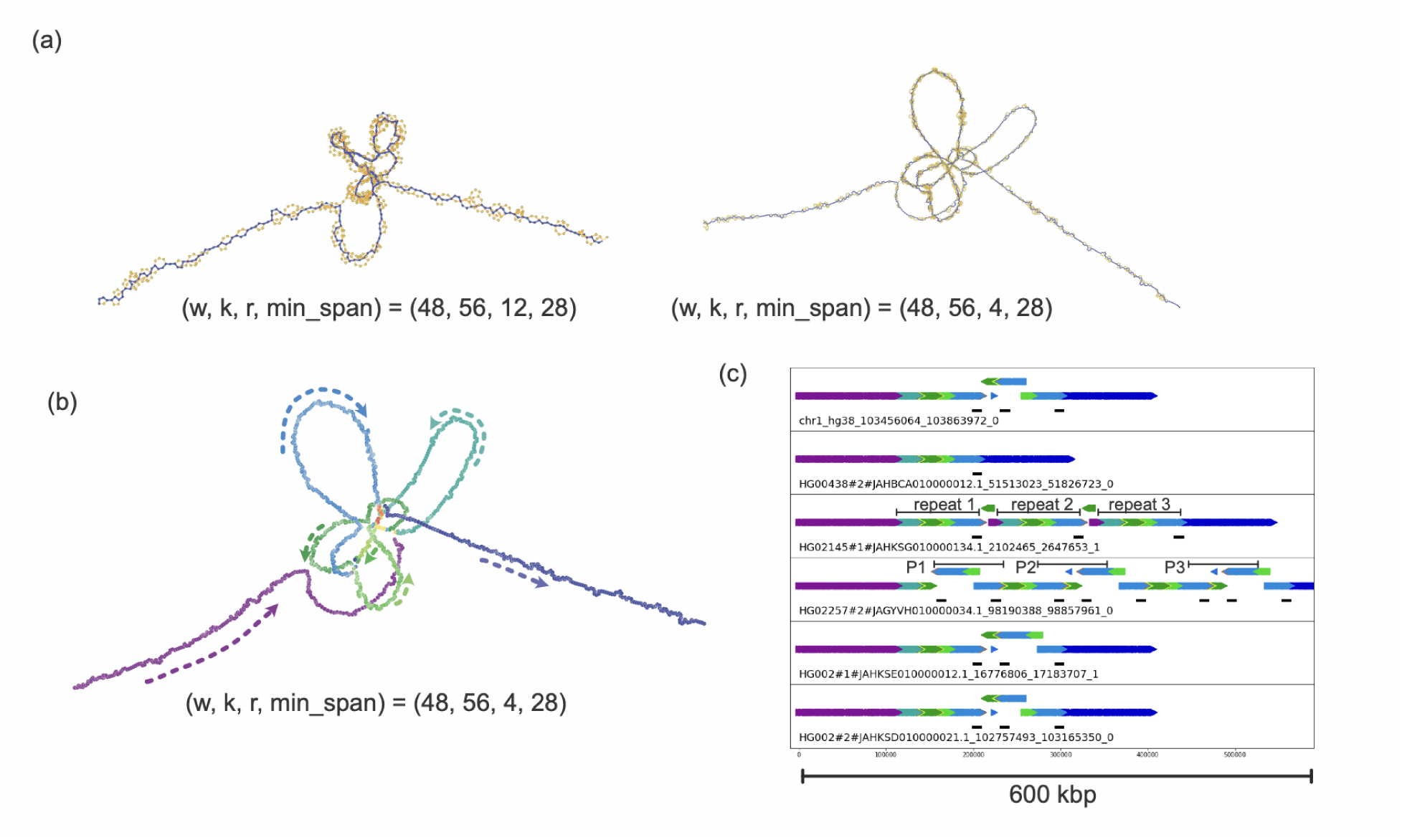
(a) AMY1A MAP graph in two different scales (b) An example of the MAG graph principal bundle decomposition for visualizing the repeat structure. (c) The principle bundle plot of 6 selected haplotypes. The AMY1A genes are annotated as the black track underneath the main bundles. The repeat regions in HG02145#1 are highlighted with repeat1/2/3 and the palindrome repeats in HG02257 are annotated with P1, P2, and P3.

In PGR-TK, we provide tools analyzing a MAP graph to “re-linearize” the graph into a set of “principal bundles.” We design the algorithm to generate the principal bundles representing those consensus paths which are most likely corresponding to repeat units in the pangenomes. The algorithm searches the paths that the majority of the pangenome sequences go through without branching as the principal bundles. This is analogous to identifying the contigs^51,52^ in genome assembly algorithms.

**Figure 2b** shows the principal bundle decomposition of the AMY1A region MAP graph. The major part of the graph is partitioned into different principal bundles which are shown in different colors. When tracing each sequence through the graph, the path of the sequence overlaps with the principal bundles. By identifying the overlaps on the principal bundles, we can see which tentative repeat units are inside the sequences of interests. **Figure 2c** shows the tracing results of 7 selected genomes over the AMY1A region.

With a linear representation derived from the MAP graph, we can identify and potentially classify the repeat structures. In the seven genomes selected for the **Figure 2c**, HG00438#2 mostly follows the principal bundles without repetition. The GRCh38 one has one relatively simple invert repeat (forming a palindrome region). HG02145#1 has mostly 3 copies of non-inverted repeats (labeled as repeat 1,2, and 3). HG02257#2 has 3 palindromic repeats (labeled as P1, P2, and P3). The two haplotypes from HG002 have similar structure to the GRCh38 except there is an inversion in the middle of the palindrome repeats in one of the two haplotypes. A full plot with all 96 repeat structures is shown in the **Supplementary Figure 2b**. We propose that researchers can utilize such decomposition to classify the different repeat structures efficiently for the regions they are interested in studying.

To demonstrate the importance of constructing MAP graphs with different scales, we generate the principal bundle decomposition for the big nested inverted repetitive regions between SMN1 and SMN2 for the assembled haplotypes that covered the region. As the repeat units are larger than those of AMY1A, we must adjust parameters to catch the larger structures more effectively. The repeat units in the SMN1/SMN2 region are typically 100kbp to 200kbp. As it is larger than those around AMY1A, we use (w, k, r, min_span) = (80, 56, 6, 28) for SMN1/SMN2 repeat region (**Supplementary Figure 3a, 3b**) instead of (w, k, r, min_span) = (48, 56, 4, 28) for AMY1A.

The MAP graphs also enable visualization of the much smaller 5kbp KIV-2 tandem repeats in LPA^53,54^. To capture the structure, we need a higher resolution. We find that we can resolve the repeat structure of the LPA KIV-2 tandem repeats with parameters (w, k, r, min_span) = (56, 56, 1, 0). With such parameters, PGR-TK generates a set of dense minimizer anchors that are suitable for distinguishing the small differences between repeat copies, including single base changes (See **Supplementary Figure 4**).

### Visualizing and Analyzing the Highly Polymorphism HLA Class 2 Locus

Major histocompatibility complex (MHC) region in human genomes is highly polymorphic. The genomic sequences of MHC are fundamental for understanding a human’s adaptive immune system and autoimmune diseases^55,56^. Unfortunately, due to the complexity and the polymorphic nature, it has been challenging to get a complete picture of the MHC genomics in the human population and to benchmark variant calling in the most variable regions^39,57^. The HPRC assemblies provide a new opportunity to analyze the MHC sequences with nearly fully assembled sequences of the region.

To demonstrate how PGR-TK can help to analyze MHC, we focus on the HLA class II locus (hg19, chr6:32,130,918-32,959,917). We fetch the HPRC HLA class II haplotype sequence by anchoring them with more conserved flanking regions. As the PGR-TK does not use sequence alignment to construct a pangenome graph, we can construct the MAP graphs of the MHC regions in just several minutes, and the principal bundle analysis can help to understand large scale structural organization of the HLA class II region.

The MAP graphs of 5 selected haplotypes are shown in **Figure 3a**, with the blue paths highlighting five haplotypes in the full pangenome graph from the HPRC assemblies. The topology of the graphs is more complicated than most other regions in the human genome due to the high polymorphism of the regions. In fact, many haplotype sequences do not align to the primary contigs in the reference genome (hg19 or GRCh38) as the sequences are simply distinct, and the new human genome assemblies reveal many new fully assembled MHC haplotypes, in contrast to most of the GRCh38 alternate loci that are only partial MHC sequences.

**Figure 3.**
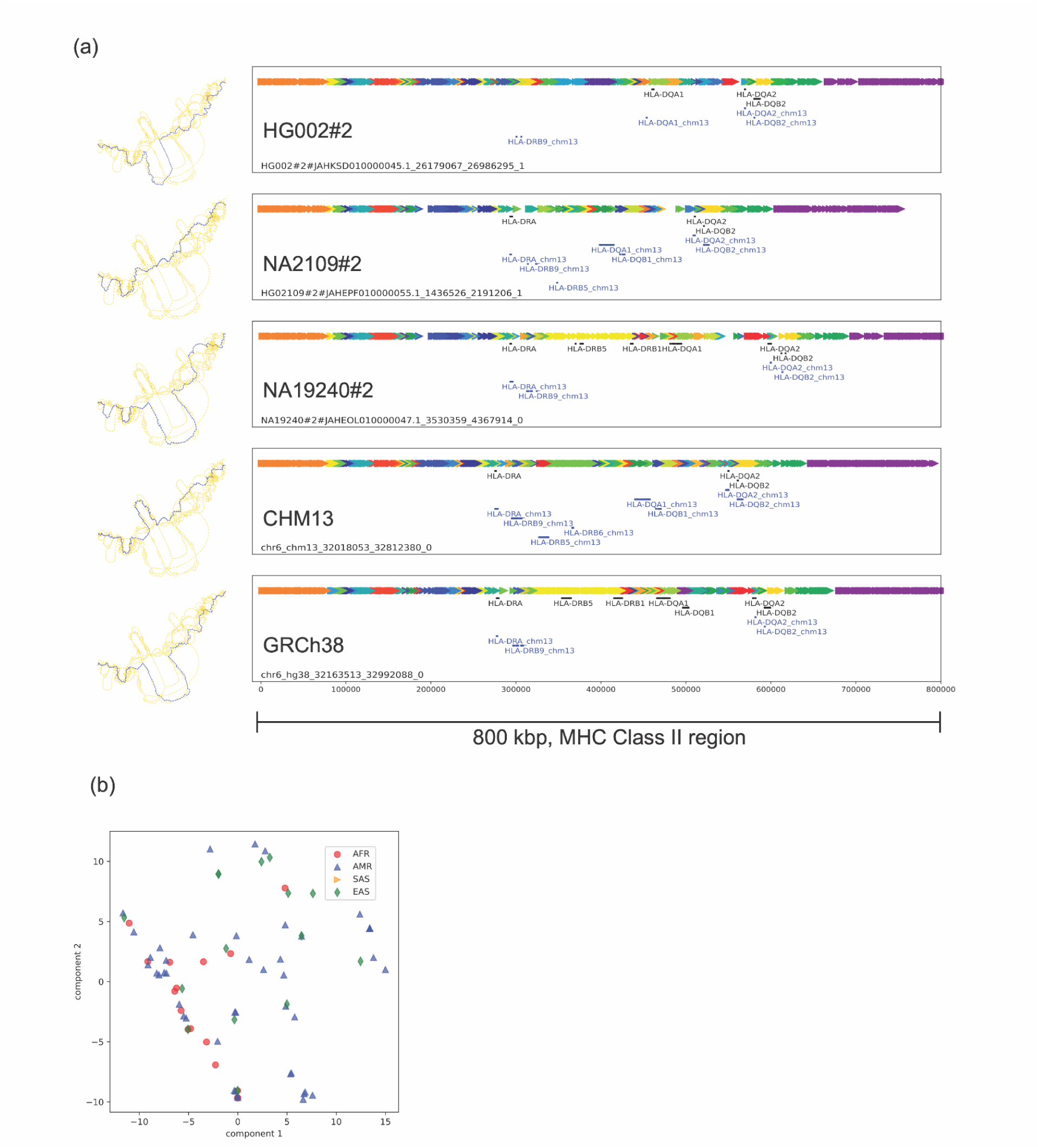
MHC Class II region MAP graph: (a) different haplotypes (highlighted in blue) in the highly polymorphic region with parameters (w, k, r, min_span) = (128, 56, 12, 8) and the principal bundle decomposition of each haplotypes. The gene regions from GRCh38 and CHM13 are fetched and shown as additional tracks if homology is found in those haplotypes. (black tracks: HLA-DR/DQ/DB from GRCh38, blue tracks: HLA-DR/DQ/DB from CHM13. (b) PCA plot of the pangenome sequencing corresponding to the MHC Class II region.

**Figure 3a** also shows the principal bundle decomposition of the selected haplotypes and we map the HLA class II HLA-DRA, HLA-DRB, HLA-DQA, and HLA-DQB gene families found in GRCh38 and CHM13 on top of each haplotype. The diversity between the haplotypes is high in the region containing the genes between the two flanking segments, but, qualitatively, common principal bundles are arranged similarly on the flanking 5’-end (e.g., the orange, cyan, red and blue segments on the left) and flanking 3’-end (e.g., yellow, darker green, and purple segments on the right). In the middle, the structure of the haplotypes can be highly dissimilar from each other, which is why some genes are only contained in alternate loci for hg19 and GRCh38. As the haplotype sequences are quite distinct, we do not find all the HLA-DRA, HLA-DRB, HLA-DQA, and HLA-DQB gene family sequences previously annotated on GRCh38 or CHM13 mapped on the selected haplotypes with a quick mapping method utilizing the SHIMMER index. This does not imply those genes are missing but the exact DNA sequences including the non-coding part of the genes might be quite distinct and have many structural variations between the different individuals.

**Supplementary Figure 5a** shows an extended principal bundle analysis plot including many other haplotypes from the HPRC pangenome. In order to see if there are overall distinct clusters of the HPRC HLA class II haplotypes, we construct a binary vector by projecting it to the principal bundle and perform a principal component analysis using such vectors of all sequences. We do not observe a clear clustering structure that is correlated with the known ethnicity groups of the samples (**Figure 3b**). It implies more additional haplotype sequences will be necessary to reveal the population structure of this region, and additional samples from future HPRC assembly releases will help reveal additional structures.

Additional work will be needed to provide more quantitative analysis for the organization of the HLA class II. The MAP graph could provide a foundation for the future related work to fetch the related sequences for detailed sequence comparisons and gene annotations.

### Survey on the Genome in Bottle Challenging Clinical and Medically Relevant Genes with the MAP graphs

The Genome in the Bottle Consortium provides variant call benchmarks on seven benchmark genomes^40,58,59^. These benchmarks were initially formed by integrating multiple short-read technologies, but the latest version integrated linked-read and long-read technologies to form benchmarks in regions difficult to map with short reads. However, a set of 395 challenging but medically relevant genes were identified as substantially (>10%) excluded from the mapping-based benchmark due to long repeats, large structural variations, segmental duplications, and/or high polymorphism between the benchmark genome HG002 and the reference genome hg19 or GRCh38. A long-read genome assembly approach provided a reliable benchmark call set of 273 out of the 395 genes ^40^. The remaining 122 were still excluded mainly because they were not accurately assembled or no benchmarking tools exist to compare different representations of complex variants in the genes. Overall 395 genes have recently shown to include high levels of polymorphism across different ethnicities making it highly challenging to represent their variations^60^.

The current GIAB variant benchmarks focus on a small set of seven well-characterized genomes with extensive short-, linked-and long-read data to ensure robust benchmarks and more tractable method development and benchmark evaluations. Meanwhile, limited representation of genomes in a population may miss significant structural variants, additional copies of genes, and context for important variants for diseases not observed in a smaller dataset. Given that increasingly accurate long-read and assembly level data are being produced at pangenome scales now, we are surveying how we can utilize such resources to benchmark variant accuracy at a broader population level for the challenging clinical and medically important genes. Such pangenome analysis will help to generate guidelines for future practice.

With PGR-TK, we extract all sequences from the HPRC year one release (94 haplotype assemblies), and CHM13 v1.1, GRCh38 and hg19 of all 385 CMRG. We generate a MAP graph of each gene and output in GFA format (see **Supplementary Material**). For each graph, we derived two metrics to estimate (1) the degree of polymorphism among the pangenomes, and (2) the repeat content taking account of the variations of the pangenomes.

To estimate the degree of polymorphism, we consider a diffusion process in the graph (see **Methods**) and derive an entropy-like quantity from a normalized equilibrium distribution (see **Supplementary Material** for the table of the metrics of each gene). The higher entropy values indicate more complicated graphs likely from the polymorphism from different genomes. The diffusion weight on each vertex is also associated with multiplicity and repetitiveness of the corresponding segments in the pangenomes. We pick the average of the top 32 diffusion weights from the MAP graph of each gene as a simple metric to measure the most challenging repeat content within a gene.

To gain insight about the challenge for calling variants of the CMRG set at a pangenome scale, we plot the diffusion entropy versus the maximum local repeat weights for each gene (**Figure 4a**). As there are no obvious correlations, these two quantities provide independent measurements of two aspects of the MAP graph structures of these genes. We find, unsurprisingly, high repetitive genes are harder to create a reliable variant benchmark call set.

**Figure 4:**
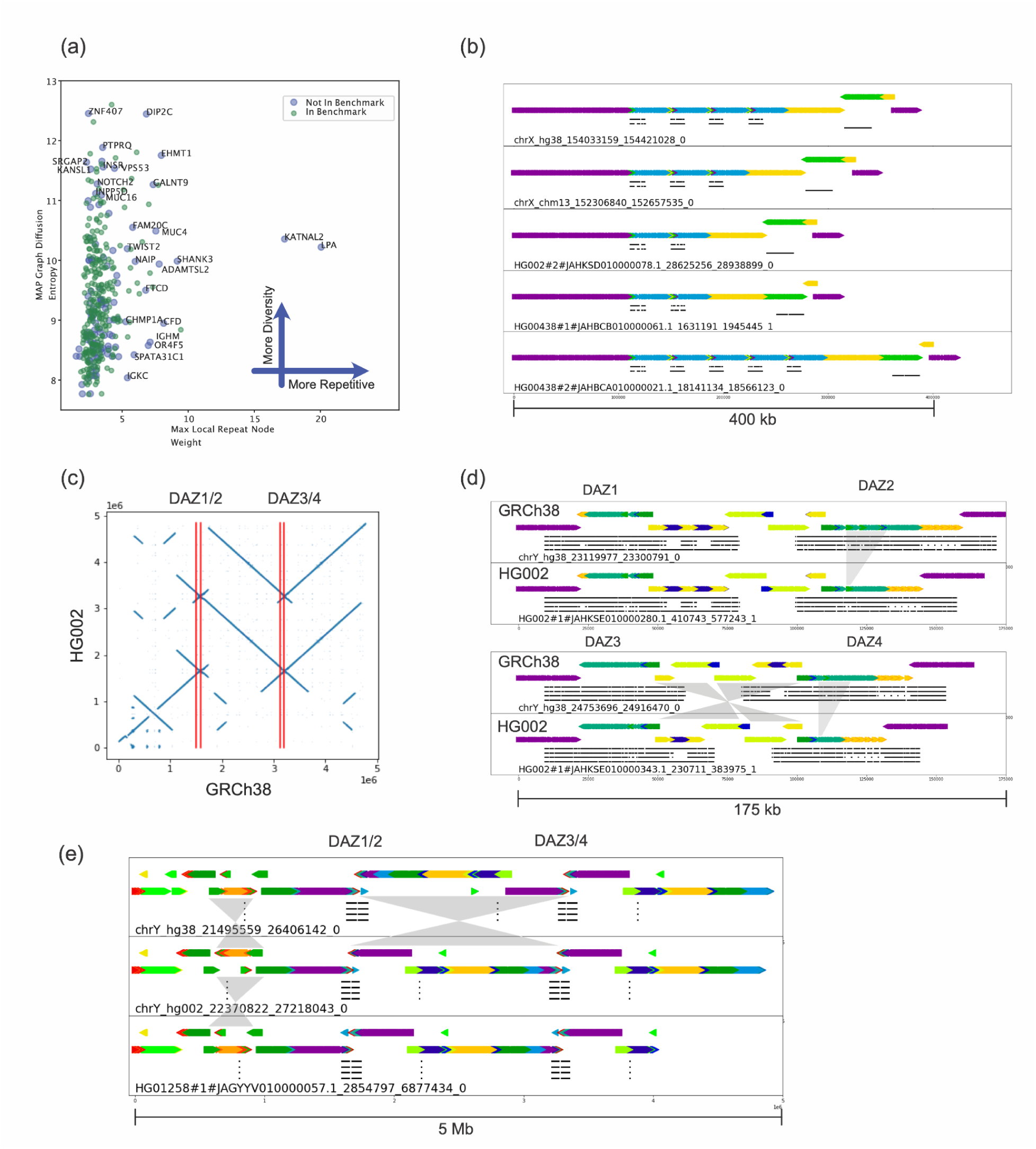
(a) MAP graph diffusion entropy vs. repetitiveness survey for the 385 GIAB challenge CMRGs. (b) MAP graph principle bundle decomposition shows the repeat number changes of the OPN1LW, OPN1MW1/2/3 to FLNA loci. Auxiliary tracks: top OPN1LW, middle OPN1MW1/2/3, bottom FLNA. (c) Dot-plot of the HG002 assembly vs. GRCh38 over the 5Mb regions spanning the whole DAZ1/2/3/4 showing an inversion between DAZ1/2 and DAZ3/4 loci. (d) MAP-graph principal bundle decomposition for DAZ1/2/3/4 system in two separate palindrome loci. Four auxiliary tracks indicate the local matches to DAZ1/2/3/4 from the top to bottom. Compared to the GRCh38 DAZ2, the hg002 DAZ2 missed a chunk (~10kb) of the darker green segment. The intergenic region between DAZ3 and DAZ4 has a rearrangement that can be described either as an incomplete inversion of GRCh38, or two separate insertions and deletions. (e) The MAP graph principle bundle decomposition of the whole 5Mb region spanning through the whole inversion region between DAZ1/2 and DAZ3/4 loci of the HG002 T2T assembly and the HG1258 assembly from HPRC year one release.

Many highly repetitive genes are excluded from the current CMRG benchmark set. We do not observe that higher entropy is correlated with the reduction of the gene in the benchmark set. We find the high entropy genes also span larger regions in the genome. While the entropy can indicate the complexity of variations in the population, we observe different clustering structures of the top entropy genes. (Please see the comparison of the MAP graph PCA plots of SNTG2 and KMT2C in **Supplementary Figure 6**.)

We highlight several genes with high entropy or high repetitiveness. The MAP graphs and the IGV view of the pangenome assemblies of a selected set genes (LMF1, ANKRD11, SRGAP2, KMT2C, LPA, MUC4, MUC3A, KATNAL2, FLG) aligned to GRCh38 are shown in **Supplementary Figure 7**. As shown in LPA (associated with Coronary Disease^61^, **Supplementary Figure 4**), KATNAL2 ( loss of function variant discovered in autistic proband^62,63^) also has long tandem repeat variations in the HPRC pangenome cohort. We find the number of the 5.8kbp repeats inside KATNAL2 ranges from 3 to 25 (**Supplementary Figure 8**). Applying MAP graph decomposition on genes such as KATNAL2 with a big pangenome reference panel will provide additional insights to the natural of the variability of the repetitiveness and its impacts to the underlying biology like other more well studied gene, e.g. LPA, in the coming years.

GIAB is using the fully assembled HG002 chrX and chrY from T2T to form new small variant and structural variant benchmarks. The assembly fully resolves the medically relevant ampliconic genes^41,64,65^ OPN1LW/OPN1MW/OPN1MW2/OPN1MW3 and DAZ1/DAZ2/DAZ3/DAZ4, but the variation in these genes is too complex for current approaches to make reliable variant calls compatible with current benchmarking tools.

For example, the genes OPN1MW and OPN1MW2 are inside a 74 kb deletion in HG002 relative to GRCh38, so HG002 contains only 2 of the 4 copies of the array in GRCh38 - OPN1LW and one copy of OPN1MW/OPN1MW2/OPN1MW3. Dipcall^66^ can call the 74 kb deletion and variants in the other gene copies, but it may be possible to align the assembly to GRCh38 in alternative ways. The visualization from PGR-TK makes clear the varying number of genes in this array in each haplotype in **Figure 4b**, which is important for some phenotypes like color blindness, since seeing full color requires OPN1LW and at least one copy of OPN1MW/OPN1MW2/OPN1MW3^64,67^.

Another important gene family DAZ1/DAZ2/DAZ3/DAZ4 are in a set of nested palindromic repeats. It has been reported that partial deletions in this region may cause male infertility^65^. It would be useful to understand the natural distribution of non-pathogenic structural variants across this ampliconic gene cluster. DAZ1 and DAZ2 are ~1.5 Mbp from DAZ3 and DAZ4, and HG002 has a 1 to 2 Mbp inversion relative to GRCh38 with breakpoints in the segmental duplications that contain the DAZ genes (**Figure 4d,4e**). In addition to the large inversion, the DAZ genes contain structural variants, including a ~10kb deletion in DAZ2, 2 deletions in DAZ4, and 2 insertions in DAZ3 of sequences that are only in DAZ1 and DAZ4 in GRCh38 (**Figure 4d,4e**). PGR-TK’s ability to color and visualize variation with the principal bundle decomposition algorithm at multiple scales enables intuitive understanding of this type of very complex variation, which would be very difficult to represent and understand in VCF.

## Discussion

With the advance in DNA sequencing technologies, more comprehensive human genomes at, or close to, telomere to telomere will be collected and made available in the coming years. It will enable researchers to study and characterize those previously inaccessible complex, but likely relevant, regions. The current Human Pangenome Reference Consortium assembly release has significantly impacted our understanding of the human genome architecture. It will also be essential for building applications for clinical and medical tests and diagnostics soon. Flexible and scalable computational tools for analyzing pangenome level genome assemblies will be part of the vital task of improving the practice of precision medicine with rich genomic data such as those from HPRC.

Many of the recently developed pangenome analysis tools allow graph analysis at the whole genome level^22–24^. Meanwhile, the richness of diversity of human genomes over the repetitive regions poses unique challenges for analysis. In our work developing PGR-TK, we focus on providing a flexible library of useful algorithms. Furthermore, it enables analyzing the genome assemblies such that a developer or a researcher can rapidly access certain complex regions by adjusting parameters for visualization and integrating with subsequent analysis.

Currently, a user needs to specify a parameter set to generate the minimizer anchors of a proper density for analysis. Higher density (smaller “w” and “r”) gives more resolution but is also more computationally intensive. On the other hand, coarse-grained views with larger “w” and “r” allow examining bigger regions at once but lose the details of the variations at smaller scales. Note that with the same “w,” and “k,” different “r” provide a hierarchical structure. Namely, the minimizer set of a choice r_1_ is a subset of the minimizer set of r_2_ if r_1_ > r_2_. In our current work, we do not utilize this interesting property yet. However, it can be leveraged to do multi-scale analysis on the fly. Here, we take a pragmatic point of view on choosing the parameter set. As the PGR-TK can do a quick regional level analysis, users can try different parameters within minutes. In the meantime, a theoretical study of the mathematical properties of how (sparse hierarchical) minimizers distribute on DNA sequences depending on their context will be necessary for future improvement and understanding of their limit^68^.

Each main building unit (the vertex) of the MAP graph represents a set of closely related sequence fragments. This is more analogous to the stringomics method proposed by Paolo^21^ than other methods building graphs on top of MSA or variant calls. Such approaches combined with the sparse minimizers is efficient to reduce the computation complexity (fewer vertices) to represent larger-scale scale structures. Complementary to that, PGR-TK provides an interface to fetch the sequences within each vertex such that it is possible to combine a MAP graph with other graph analysis approaches, e.g. Cactus graph^25^ and A-de Bruijn Graph^36^ for base level analysis, e.g. variant calling, genotyping, and point mutation analysis, with recursive hybrid graph data structures.

In the Results, we demonstrate how to use the PGR-TK for studying and characterizing repetitive regions (AMY1A, LPA, KATNAL2, etc.) and the highly polymorphic HLA Class Il region. We also derive two metrics for measuring the polymorphism and repetitiveness in human pangenome to more systematically survey complexity of a large set of medically and clinically relevant genes. We present a tool in PGR-TK backed by a novel pangenome graph traversal algorithm, re-linearizing tangled graphs caused by repetitive sequences to principal bundles for visualization. With the principal bundle decomposition, we can automatically visualize the repetitive and non-repetitive components of haplotype assembly contig. The PGR-TK can provide intuitive qualitative information about different genome arrangement architectures with decomposition and associated visualizations. For example, it enables visualization of both the very large inversion in the DAZ locus and much smaller complex structural variation within the genes. The OPN1LW/OPN1MW gene array enables visualization of copy number of the subtly different OPN1LW and OPN1MW genes, which can affect vision, as well as nearby structural variants. In the future, we aim to extend the PGR-TK library to provide more quantitative and base-level analysis for both fundamental and translational research utilizing pangome resources.

## Methods

### Sequence Database and SHIMMER Index construction

To generate the SHIMMER index, each sequence is scanned and the symmetrical minimizers were generated with the specific minimizer window size “w” and kmer size “k”. We call this first level of minimizer. Given a reduction factor “r” > 1, additional levels of minimizer sets^33^ are generated to increase the span between the minimizers by a reduction step to facilitate pan-genomics analysis. Even with the reduction step, in some simple sequence context, e.g. long or short tandem repeats, two minimizers can remain too close to each other. A parameter “min_span” can be applied to eliminate a pair of minimizers that are too close. We use a heuristic algorithm to eliminate those minimizers that are within the distance of “min_span” to each other. This helps to reduce the minimizer density when detailed analysis for those simple context regions is not desired. Setting “min_span” to zero and “r” to one will generate the standard minimizers for each sequence.

Each pair of the reduced minimizers (SHIMMERs) are used as the key to build a hashmap to the sequence id, and coordinates and matching orientation of the minimizer pairs on the sequences.

For a large sequence set, e.g. 47 whole genome HPRC assemblies, PGR-TK utilizes the AGC format^45^ to store the sequence for efficiency. A command line tool “pgr-mdb” is developed with the PGR-TK package to create the index file on top of the AGC file. For example, for a pre-build HPRC year one assembly AGC file (1.33Gb), we create a file (`/data/pgr-tk-HGRP-y1-evaluation-set-v0_input`) include a file system path to the AGC file, `/data/gr-tk-HGRP-y1-evaluation-set-v0.agc,` and call ‘pgr-mdb’ to create the index files with a pre-specified prefix (`/data/pgr-tk-HGRP-y1-evaluation-set-v0`):

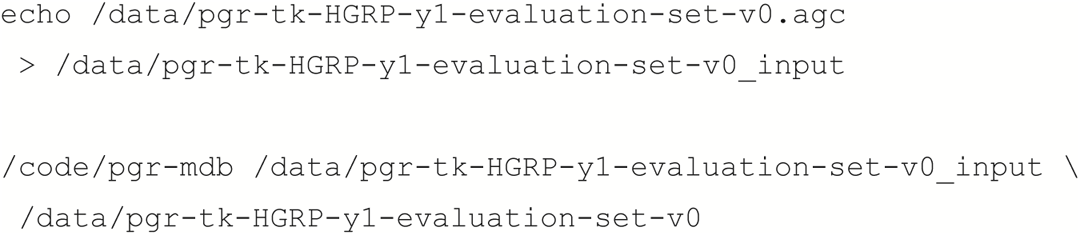

Two files will be generated in this example:

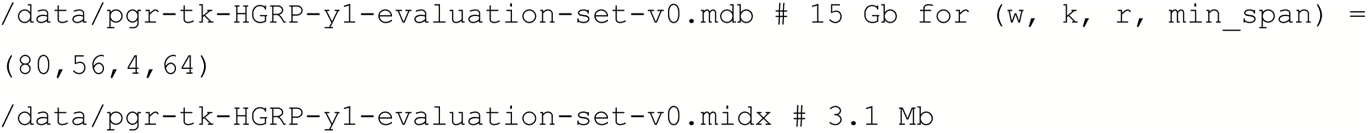

The index and sequence data can be loaded into a python workspace by

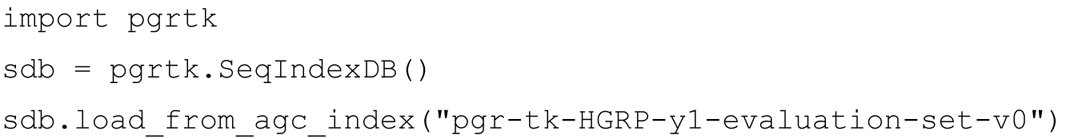

As the indexes are loaded into memory, we suggest using a computing instance which has a random access memory larger than about 4x of the index file to avoid swapping thrashing.

For smaller sequence files, the sequence database object (e.g. the “sdb” in the example above) created by pgrtk.SeqIndexDB()can create and load sequences using load_from_fastx() method. See the library documentation at https://sema4-research.github.io/pgr-tk/ for more detailed descriptions of all python objects, methods, and functions in the PGR-TK package.

### Query Sequence in the PGR-TK sequence database object

For finding homologous sequences in a PGR-TK database, we need to start with a query sequence. We can fetch a sequence in the database giving a known “data source”, “contig name” tuple and the beginning and ending coordinates. As the SHIMMERs are sparsely distributed in a sequence, the query sequence should be long enough to cover enough minimizer anchors. The python statement shows a typical code fragment to generate query results of a region of interest:

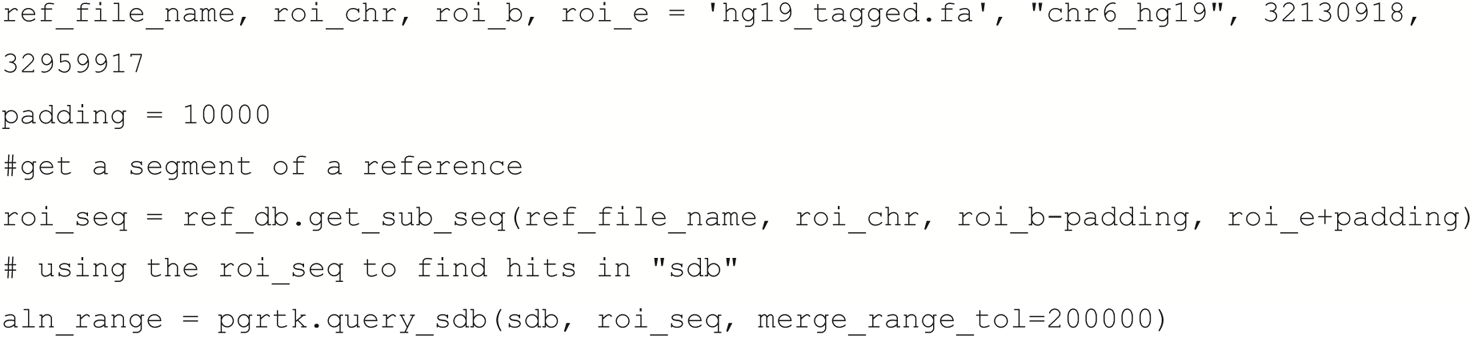

The output aln_range from the query_sdb() call contains data of the hits in the PGR-TK database. Internally, the query_sdb() method performs:

1. create SHIMMER pairs of the query sequence
2. use the SHIMMER pairs and the hashmap index to find all hits in the database
3. perform sparse dynamic programming to find sparse alignments between the query sequence and all hits in the database
4. merge the alignment segments if any of them are within the merge_range_tol parameter.

The parameter merge_range_tol is introduced to avoid alignment fragmentation when the query sequence contains a region of high polymorphism but we still want to fetch those diverse sequences for constructing the pan-genomics graph.

Typically, a user needs to process the data in aln_range for different analysis. Our example Jupyter Notebooks provides various examples for processing the output to generate dot-plot or MAP graphs, etc. (see **Supplementary Material**)

### Generate Minimizer Anchored Pan-genomics Graph (MAP graph)

The MAP graph is constructed by scanning through each sequence in the database. The vertices are simply the set of the tuples of neighboring minimizers (minimizer anchored segments). The edges are constructed by connecting minimizer anchored segments as a bi-directed graph. One can consider this as an extension of the string graph^52^, where the overlaps are the minimizers at both ends. However, in the pangenome graph, each vertex includes a set of sequence segments from multiple genomes rather than one sequence.

As the MAP graph can be constructed by scanning the SHIMMER pairs through the sequences. For a given set of n sequences S = {s_i_ | i = 0..n-1}, the vertices of the MAP graph are

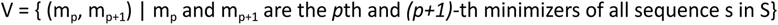

We can assign a weight w_i_ of a vertex v_i_=(m_a_, m_b_) as the total number of observed (m_a_, m_b_)-pairs in S.

The edges of the MAP graph are

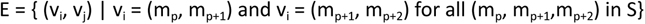

We provide a method SeqIndexDB.get_shmmr_pair_list() to get all SHIMMER pairs in a PGR-TK database. With that, we can generate a MAP graph as with the python NetworkX ^52,69^ library with such python code snippet:

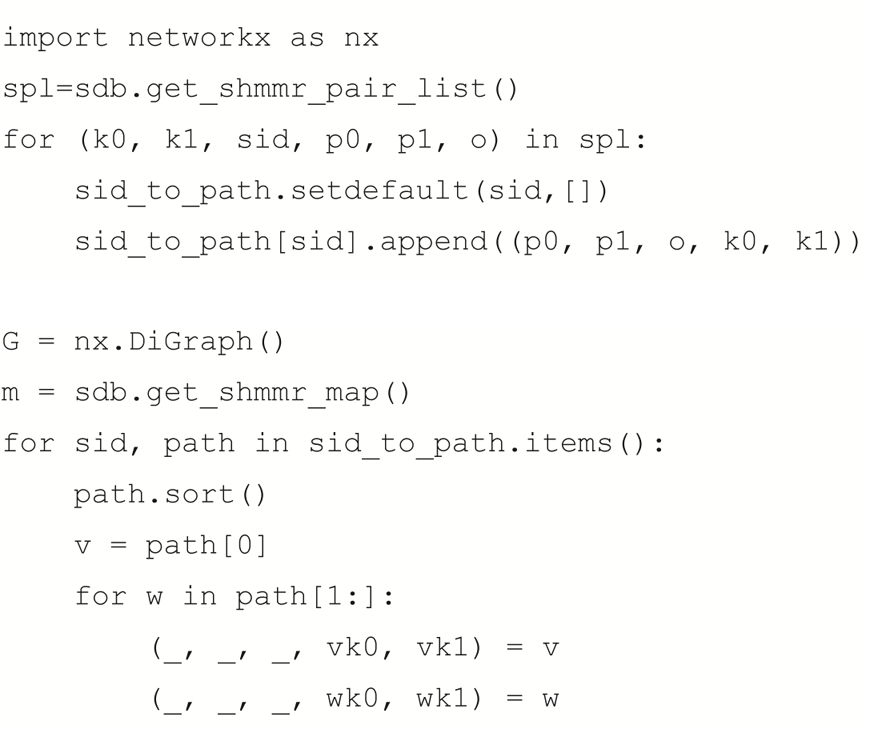

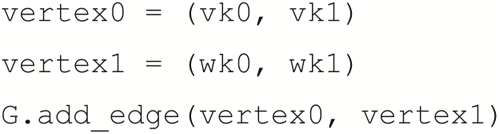

It is particularly useful when we want to do graph layout with graphviz or Gephi.

In a python session, we can use the SeqIndexDB.load_from_seq_list() to load a set of sequences of interest into the memory directly. Once the sequences are loaded, a method named generate_mapg_gfa()can save the MAP graph in a GFA format. As a standard GFA v.1 format cannot encode multiple sequences inside a GFA sequence. PGR-TK provides another function write_mapg_idx() to save the metadata that is necessary to fully reconstruct the underlying sequences in the graph and the corresponding segments to the graph vertices.

### Identify The Principal Bundles in a MAP Graph

To decompose a MAP graph into principal bundles for downstream analysis, we apply a variation of depth first search^70^ (DFS) to build the traversal trees from the graph. Our DFS prioritized vertices with high “weight” (defined as the number of sequence segments contained in a vertex) and taking account the bi-directed nature of the MAP graph.

The DFS traversal through the graph is then converted to a tree structure internally. The leaf nodes in the tree are typically when the depth first searches are terminated by no out edge from a node or a bubble or a loop is found. As we prioritize the weights of the vertices during DFS, long paths are usually corresponding to the “common” paths that most sequences in the data would go through. Rarer haplotypes typically correspond to short bubble paths in the MAP graph. Thus, they can be identified as short branches in the DFS traversal tree. We use the tree to remove those vertices in the MAP graph if those are shorter than pre-specified length in the DFS tree.

After removing the vertices corresponding to the short branches, we further remove vertices in the MAP Graph that have more than three out edges after converting the MAP graph as an undirected graph. After such removal, the graph will only consist of simple paths and we output those paths as the principal bundles.

In summary, here is the sketch of the algorithm:

1. Build a DFS traversal tree with a deep first search for a given MAP graph.
2. In MAP graph, remove vertices which corresponds to nodes in short branches of the DFS traversal tree
3. Remove branching vertices in MAP graph (by considering it as an undirected graph)
4. Output the simple paths from the resulting graph as the principal bundles.

### Principal component plot for the HLA Class II locus

To generate the principal component of the pangenome HLA Class II sequences, we convert each of the haplotype sequences to a binary vector. The binary vector has the same length of the total number of vertices of all principal bundles. Let’s call these vertices V = {v_i_ | v_i_ in principle bundles, i = 0..n-1}, where n is the total number of vertices in the principal bundles. For each sequence s, we construct a binary vector w_s_ = {b_0_, b_1_, …, b_n-1_} where b_i_ = 1 if the sequence s contains the vertex v_i_, and b_i_ = 0 if not. Then, we perform the standard principal component transformation with the binary vectors of all sequences from the HLA Class II region.

### General Workflow for Analyzing a Region of Interest

Here we outline the general workflow on how to use PGR-TK to generate MAP graph and the principal bundle decomposition

1. For a region or sequence of interests, fetch the sequence from a database for query
2. Query the whole pangenome database to get initial hits which match the query sequence
3. Filter the hits to remove unwanted matches that do match a user’s analysis objectives
4. Fetch the sequences that correspond to the hits and rebuild a smaller PGR-TK database with a new set of parameters (w, k, r, min_span) that is more suitable for analyzing the specific regions depending the region size and the scale of the feature in interest
5. Optionally, we can output the fetch sequences to files if we want to use other third party tools, for example, calling variant with dipcall, create multiple sequence alignment, or building other local pangenomic graphs with minigraph, or pggb.
6. With the smaller PGR-TK database, we can generate the region-specific MAP graph in GFA formatt or graphviz dot format through the NetworkX package.
7. Generate the principal bundles (through SeqIndexDB.get_principal_bundles() method) and trace each sequence through the bundles for visualizing and studying the pangenomic haplotype structures.
8. Re-adjust the parameters (w, k, r, min_span) and repeat (5), (6) and additional analysis on the results if necessary

### Compute Graph Diffusion Entropy and Max Repetitive Weight

It would be desirable to derive quantitative measurements so we can characterize a set of large numbers of MAP graphs fast. One thing we are interested in quantifying is “how complex a graph is”. The intuition is that if a region of the genome is more polymorphic in the population, the graph will have more alternative paths, or bubbles. We like to generate a quantity as a proxy for that. For this, we borrow the idea from network science study and spectral graph theory to consider a diffusion/random walk process on a graph^71^. For a graph, we consider a set of random walkers starting at each vertex. The random walkers can drift on the graph though the edge-connection. We can consider the distribution of the random walkers in the final equilibrium state. If a graph is relatively simple, then the final distribution will be uniform (subject to minor boundary condition corrections.) On the other hand, if the weights of the vertices or topology of the graph are more complex, we would expect the final distribution of the walkers would be less uniform and reflect the complicated nature of the graph.

The final distribution of the such diffusion process can be obtained by simple matrix multiplication iteration from the adjacent matrix of a MAP graph. Given an adjacency matrix **A**, where the matrix element **A**_ij_ = number of sequence supports edge from v_i_ to v_j_. The final distribution **P** can be written as

**P** = (1/N) lim_n->\inf_ **M**^n^ **1**, where **M** = **AD**^-1^, D is the degree matrix defined as **D**_ii_ = degree of vertex v_i_ and **D**_ij_ =0 if i ≠ j, N is the total number of vertices, and **1** is a column vector in which every element is one. (When we compute **P**, we only repeat the number of multiplications N times to approximate the final distribution.)

P is a normalized column vector [p_0_, p_1_,…,p_n-1_]^T^ such that ∑_i=0..n-1_p_i_ = 1. (See **Supplementary figure 6** for an example.) The diffusion entropy used in this work is defined as S = - ∑_i=0..n-1_ p_i_ log_2_(p_i_).

To find the highly repetitive elements inside a region of interest, we look into largest elements in the unnormalized vector **NP** as a proxy of average number of repeats considering the graph structure. We pick the top 32 elements in **NP** and use the average of those as a proxy number to estimate the repetitiveness of potential repeat units inside a region of interest.

## Acknowledgments

Certain commercial equipment, instruments, or materials are identified to specify adequately experimental conditions or reported results. Such identification does not imply recommendation or endorsement by the National Institute of Standards and Technology, nor does it imply that the equipment, instruments, or materials identified are necessarily the best available for the purpose. FJS and SB are supported by NIH grants (UM1HG008898 and 1U01HG011758-01)

## Author contributions

Conceptualization and design: CSC, AK, FS, JW, JZ

Algorithm, code and documentation development of PGR-TK: CSC

Manuscript writing: CSC, AK, FS, JZ

Manuscript revision, data management and code validation: SB, FS, JW

## Supplementary Material

### Data files

The HPRC year one release sequence and pre-built index: https://giab-data.s3.amazonaws.com/PGR-TK-Files/pgr-tk-HGRP-y1-evaluation-set-v0.tar

Scripts and source data URLs for constructing the HPRC AGC file: https://github.com/Sema4-Research/pgr-tk-notebooks/tree/main/pgr-tk-sequence-source

All GFA files, fetched sequences from HPRC year one release of the 385 CMRG: https://giab-data.s3.amazonaws.com/PGR-TK-Files/CMRG_output_dir_v0.3.3.tar

Example Notebooks using PGR-TK including code making most of (the source of) the plots in this manuscript: https://github.com/Sema4-Research/pgr-tk-notebooks

Information about a docker image with pre-built PGR-TK library and Jupyter Lab Server and Usage: https://github.com/Sema4-Research/pgr-tk/blob/main/pgr-tk-workstation/Readme.md

### Code

source: https://github.com/Sema4-Research/pgr-tk

API document: https://sema4-research.github.io/pgr-tk/

pre-built binaries: https://github.com/Sema4-Research/pgr-tk/releases/download/v0.3.4/pgr-mdb https://github.com/Sema4-Research/pgr-tk/releases/download/v0.3.4/pgrtk-0.3.4-cp38-cp38-linux_x86_64.whl https://github.com/Sema4-Research/pgr-tk/releases/download/v0.3.4/pgrtk-0.3.4-cp38-cp38-manylinux_2_28_x86_64.whl

## Supplementary Figures

**Supplementary Figure 1.**
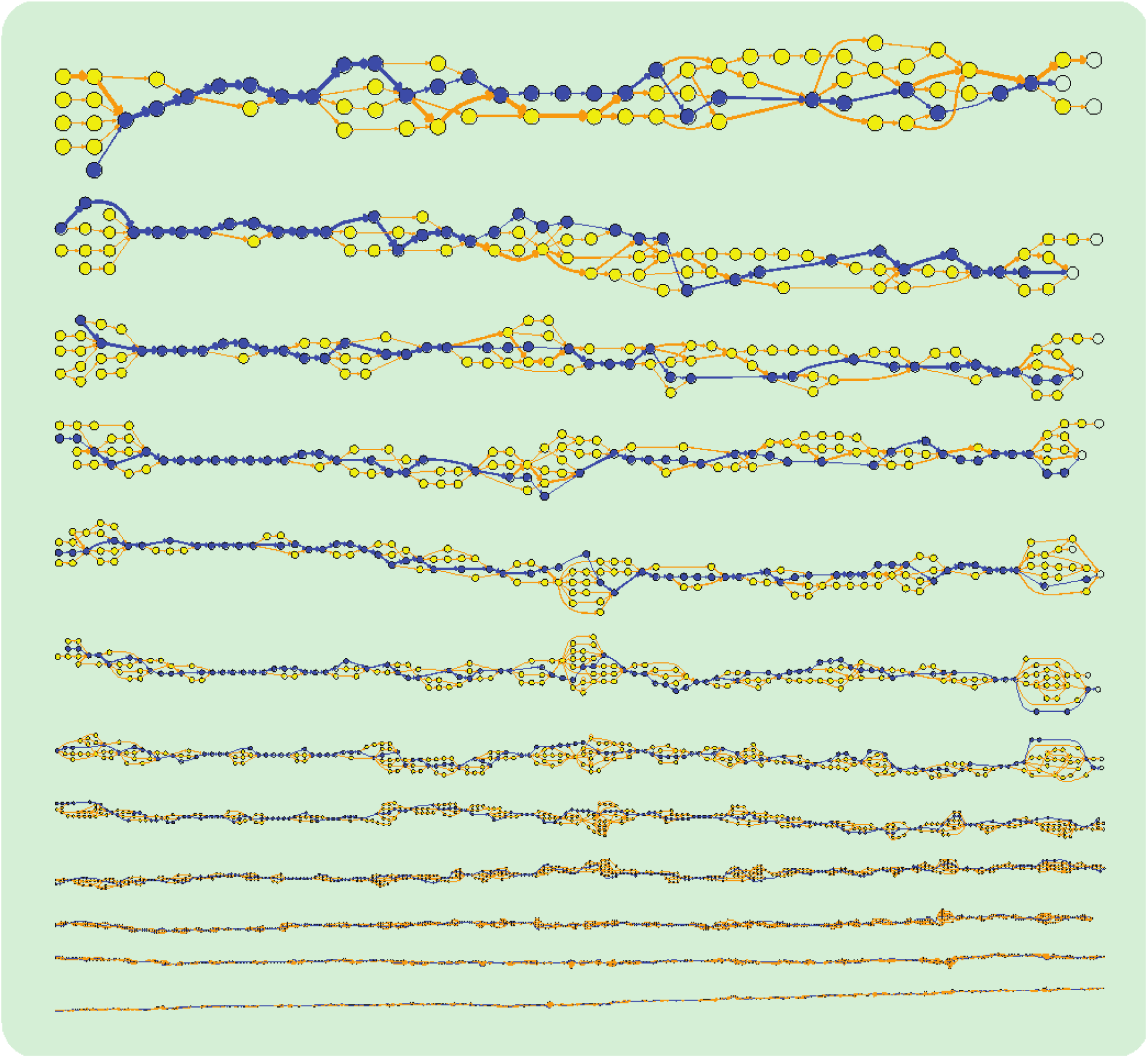
Graph of KCNE1 at various scales (Blue: Reference path, w=128, k=32, r=12 to 1)

**Supplementary Figure 2.**
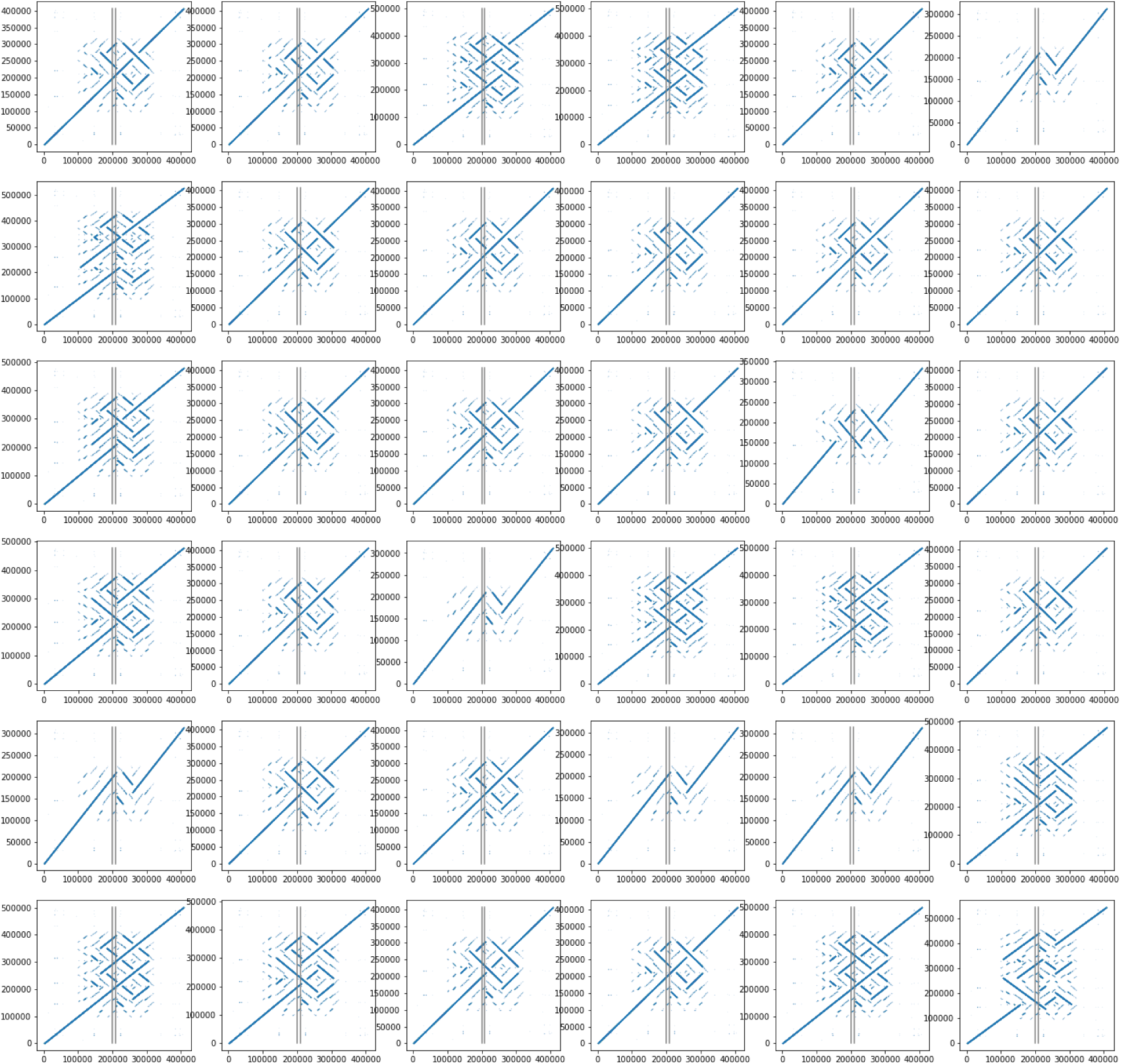
AMY1A repeat dot plots and principal bundle decomposition plots Query: hg19 chr1:104198140-104207173, parameters: {“w”: 48, “k”:56, “r”:4, “min_span”:28 } Figure 2a

**Supplementary Figure 2b:**
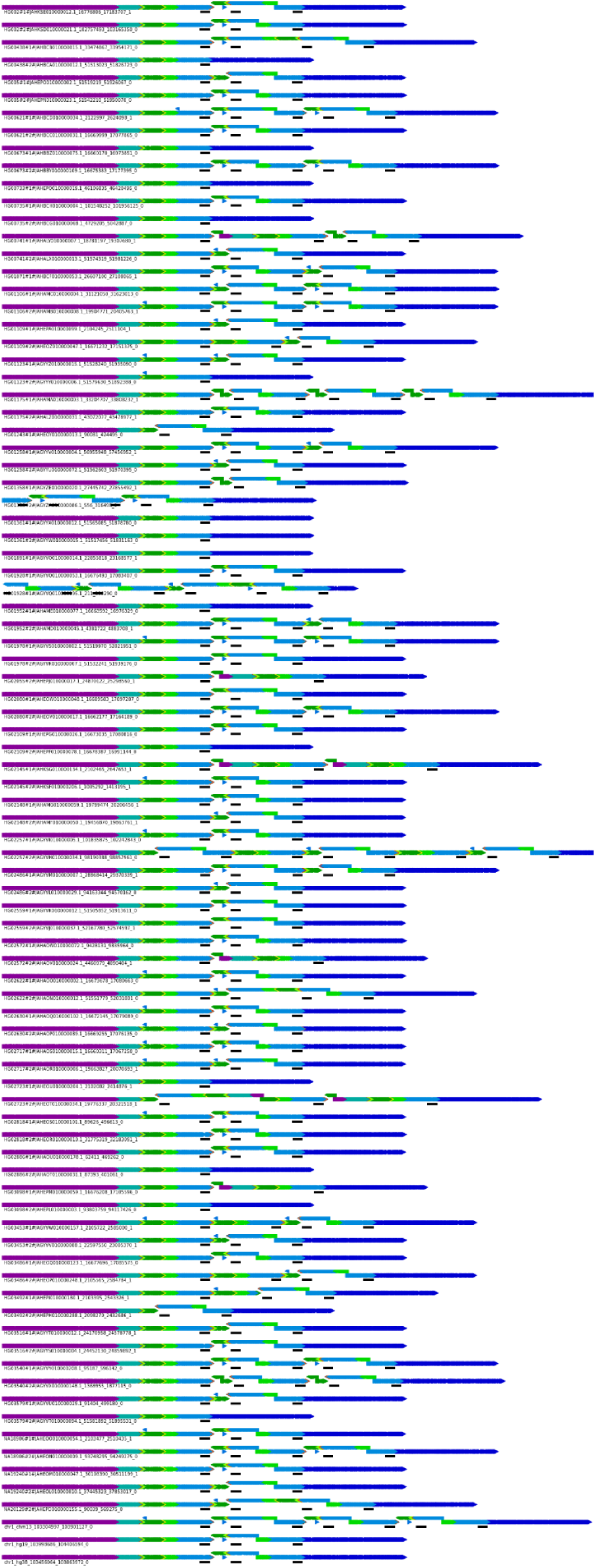
The principal bundle plots of the AMY1A repeat regions. The black short bars indicate the regions homologous to AMY1A sequence.

**Supplementary Figure 3.**
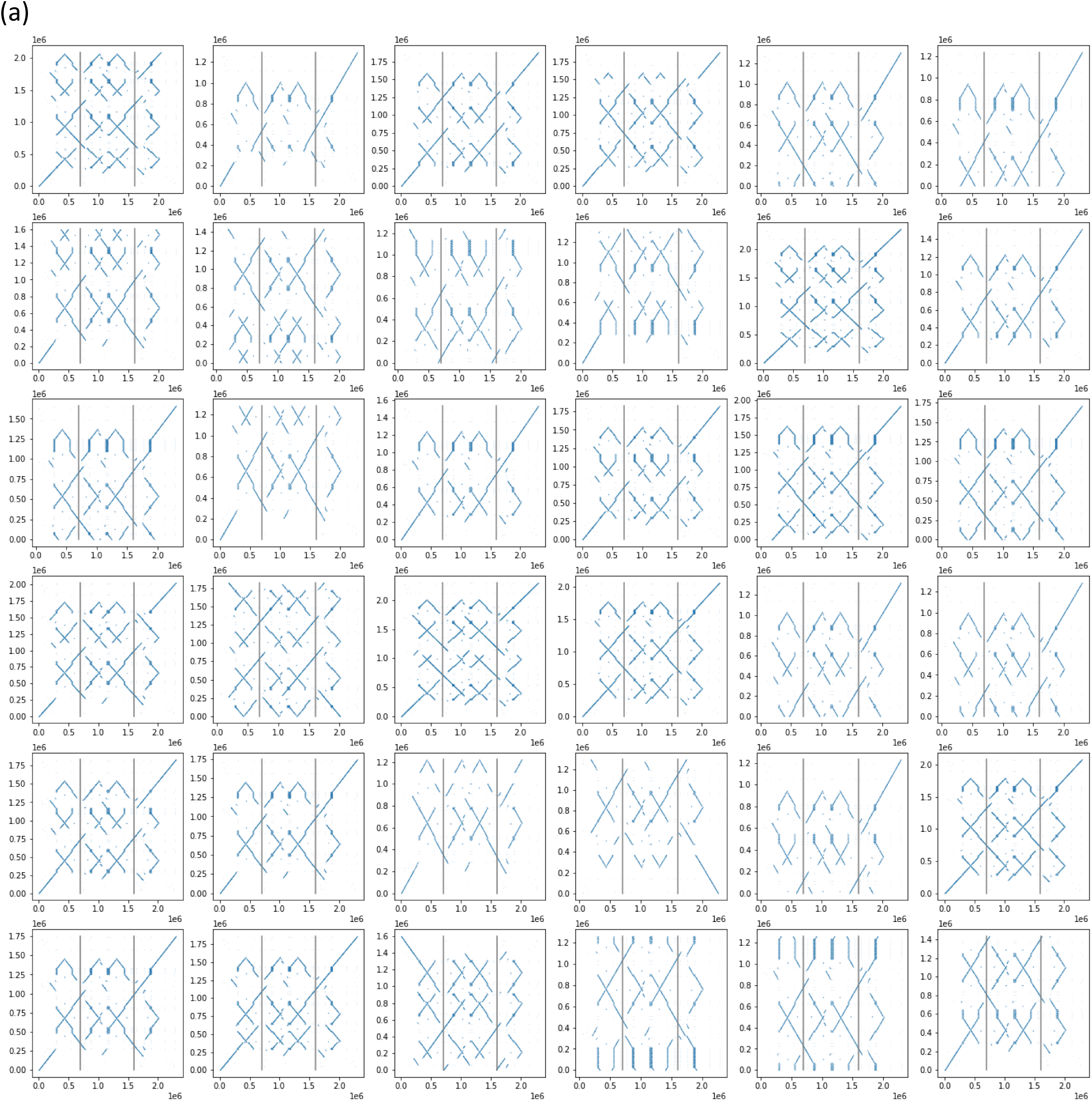
SMN1/2 repeat dot plots and principal bundle decomposition plots Query: hg19, chr5: 69345349-70248838, parameters: {“w”: 80, “k”:56, “r”:6, “min_span”:28 }. The vertical lines are the location of SMN1 and SMN2 in hg19 sequence ( x-axis)

**(b).**
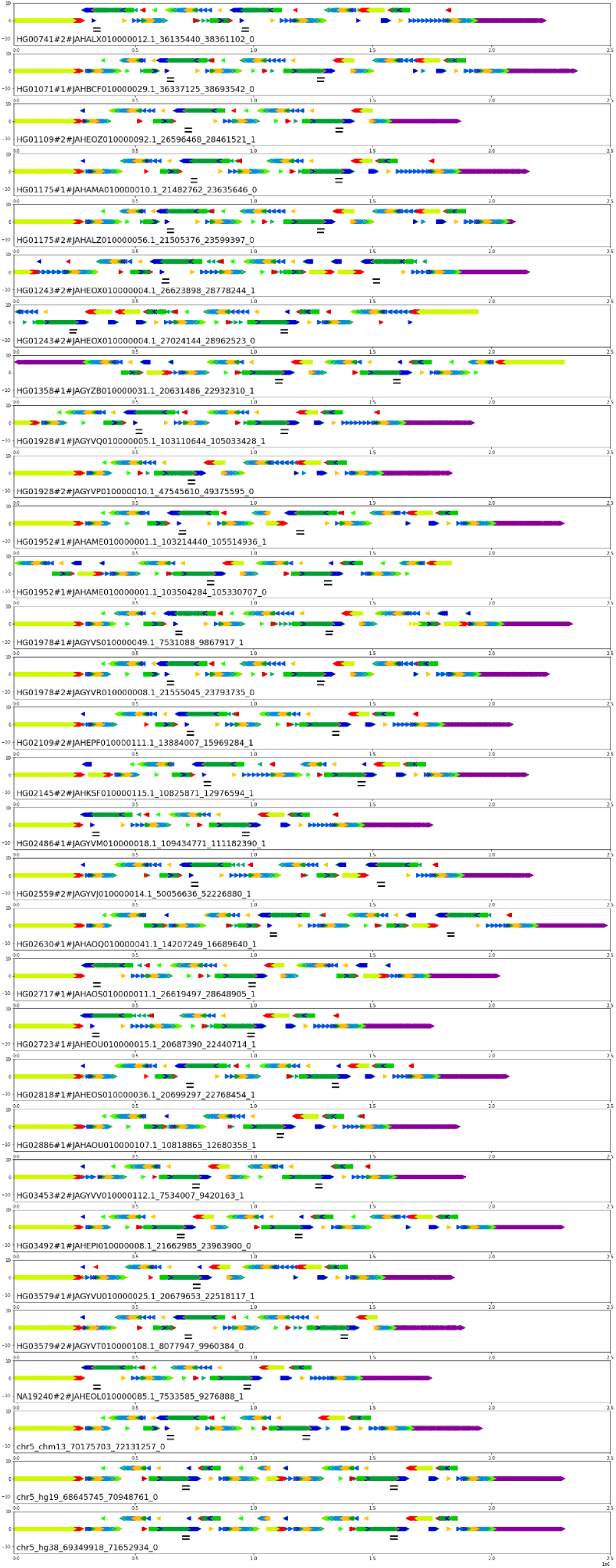
The principal bundle plots of the repeat regions between SMN1 and SMN2 of a selected set of assemblies, with the SMN1 and SMN2 gene regions labeled as black bars beneath the principal bundle plots. (Only contigs matched up more than 600kbp in this region are shown.)

**Supplementary Figure 4.**
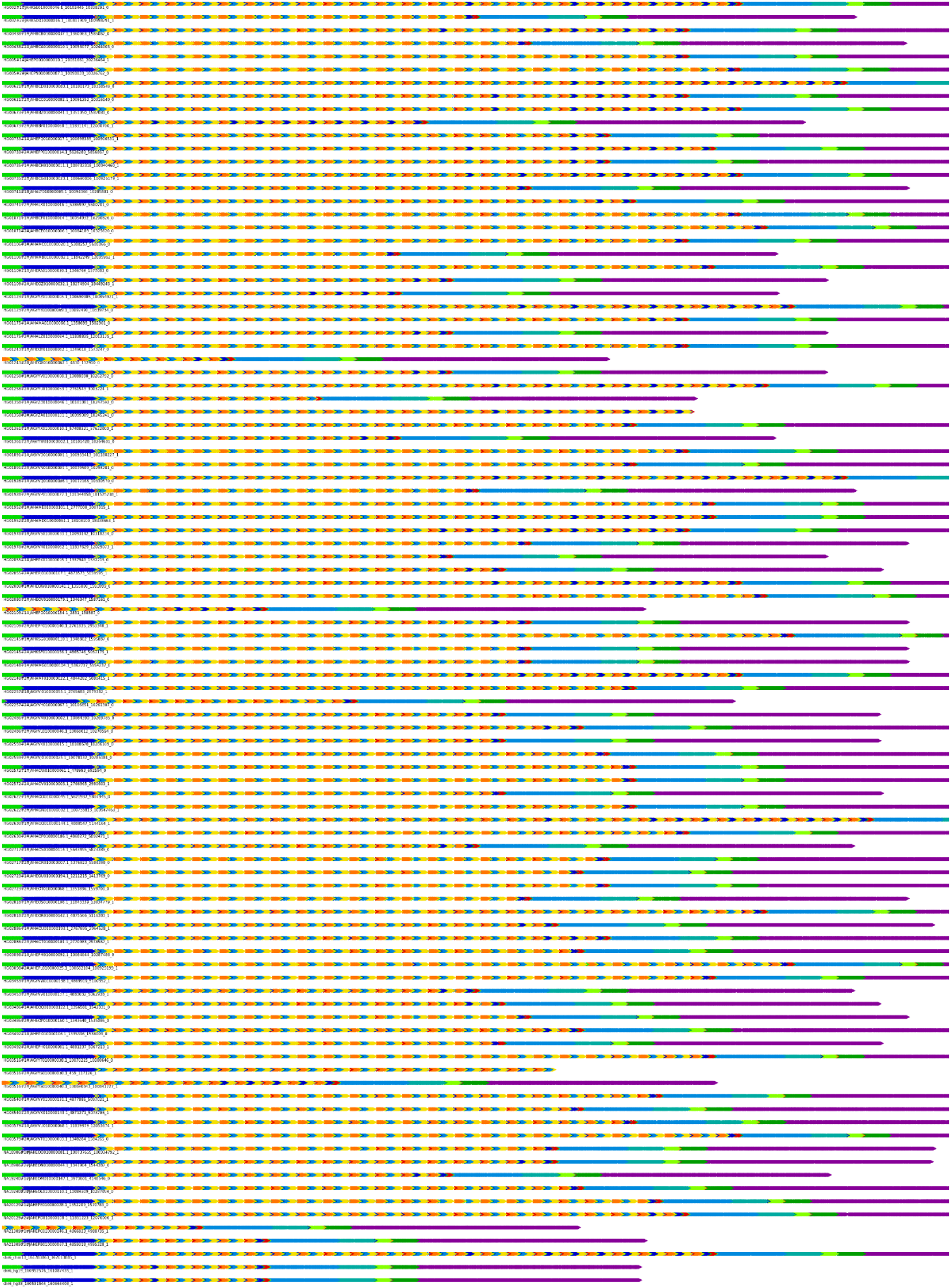
LPA, KIV-II repeats principal bundle decomposition plot Query: hg19, chr6:160952514-161087407, parameters: {“w”: 64, “k”:56, “r”:1, “min_span”:0 }

**Supplementary Figure 5.**
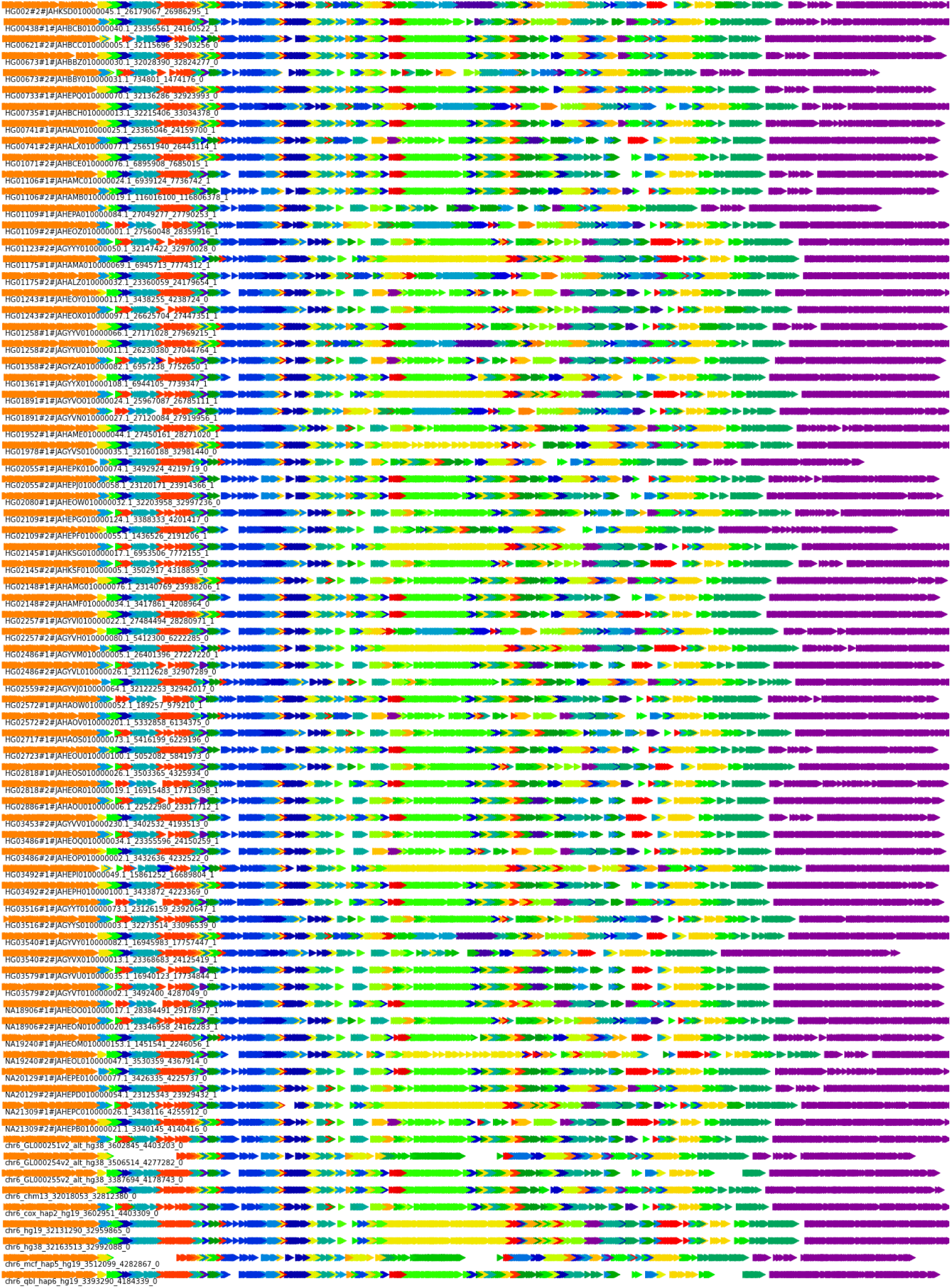
Figure 5a: HLA Class II principal bundle decomposition Query: hg19, chr6:32130918-32959917, parameters: {“w”: 128, “k”:56, “r”:6, “min_span”:28}

**Supplementary Figure 5b:**
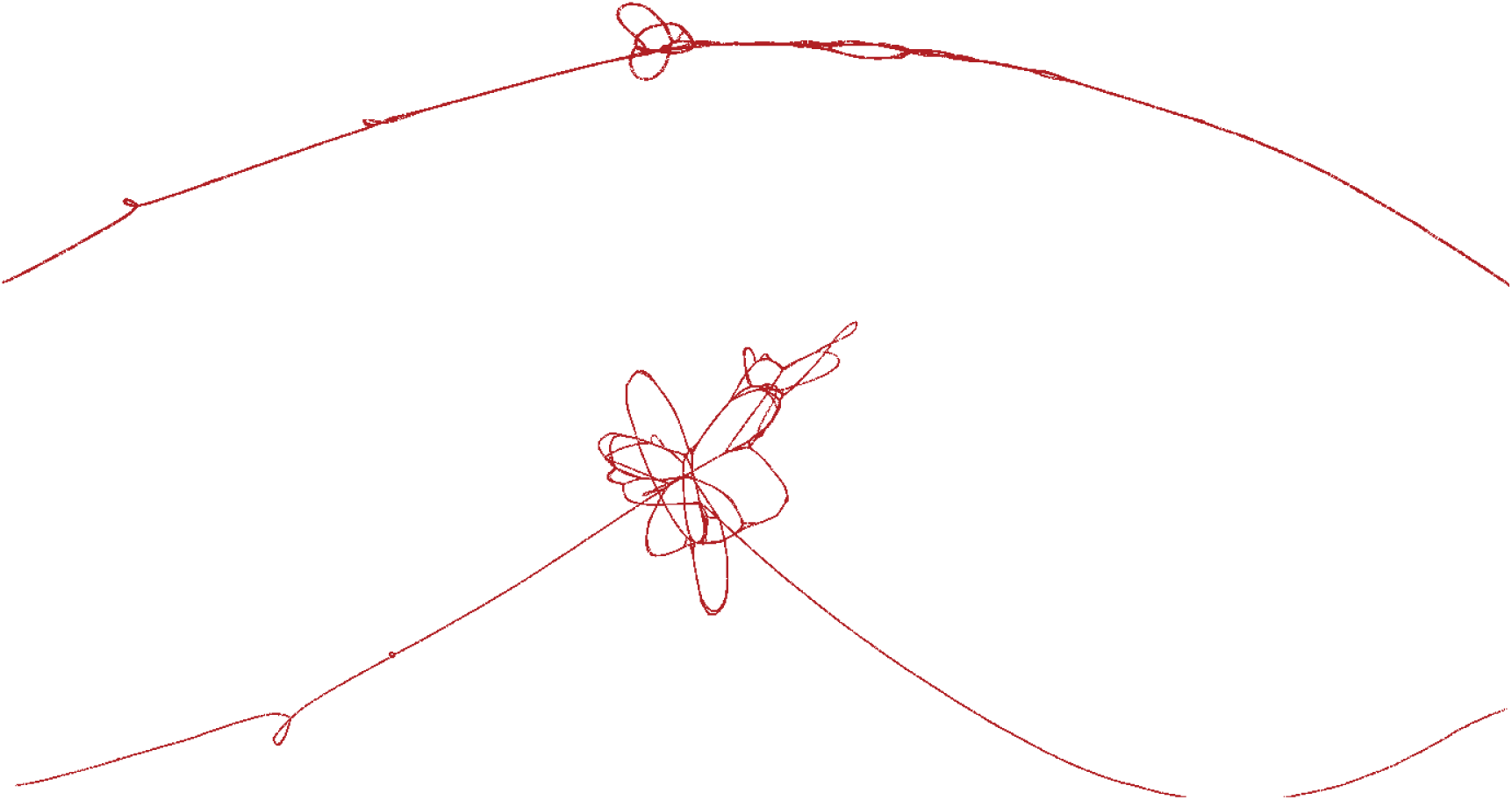
Bandage layout of the GFA files of the HLA-Class II region (hg19 chr6:32130918-32959917) in two different scale Top: (w,k,r) = (128,32,6), 4,907 Nodes, 6,395 edges. Bottom: (w,k,r) =(128,32,2), 14,620 nodes / 18,561 edges

**Supplementary Figure 6a.**
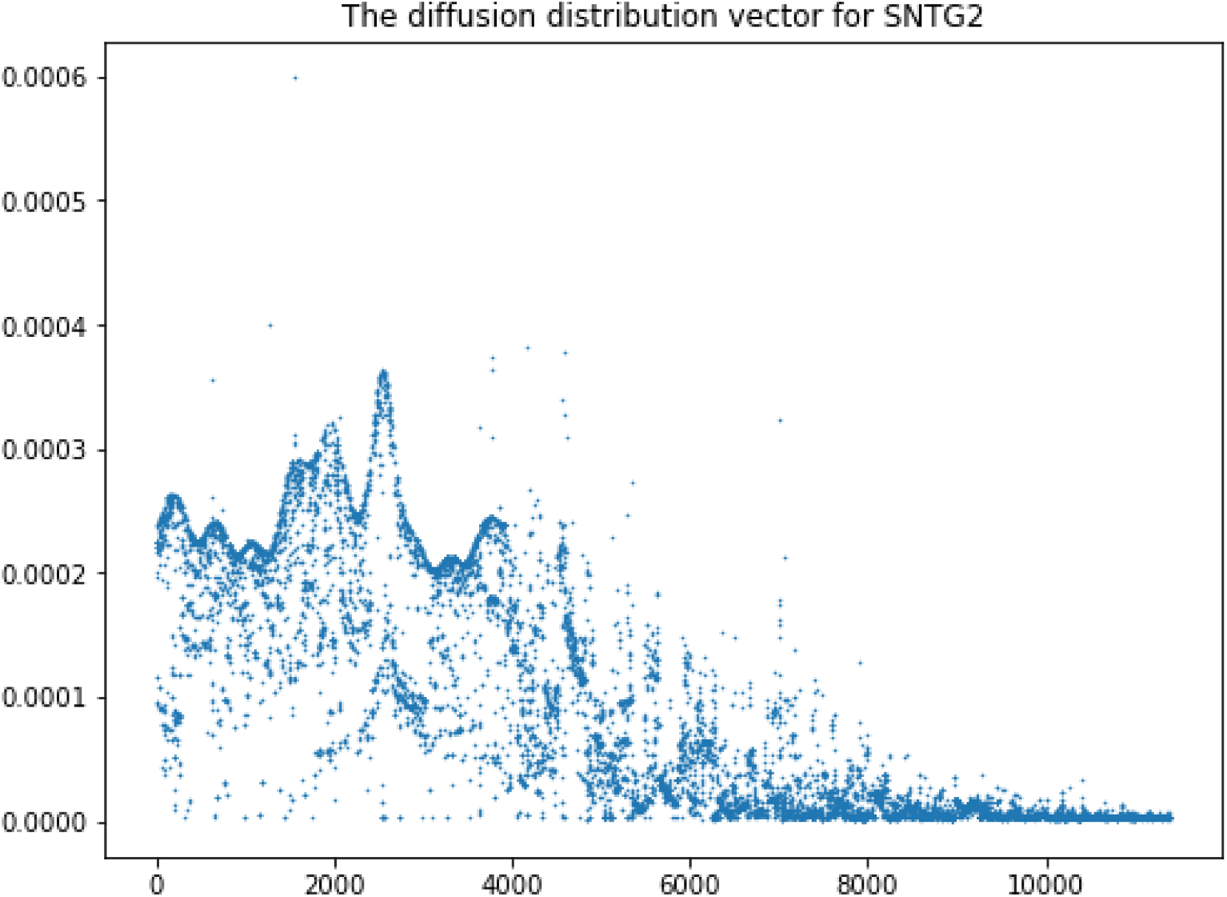

**Supplementary Figure 6b:**
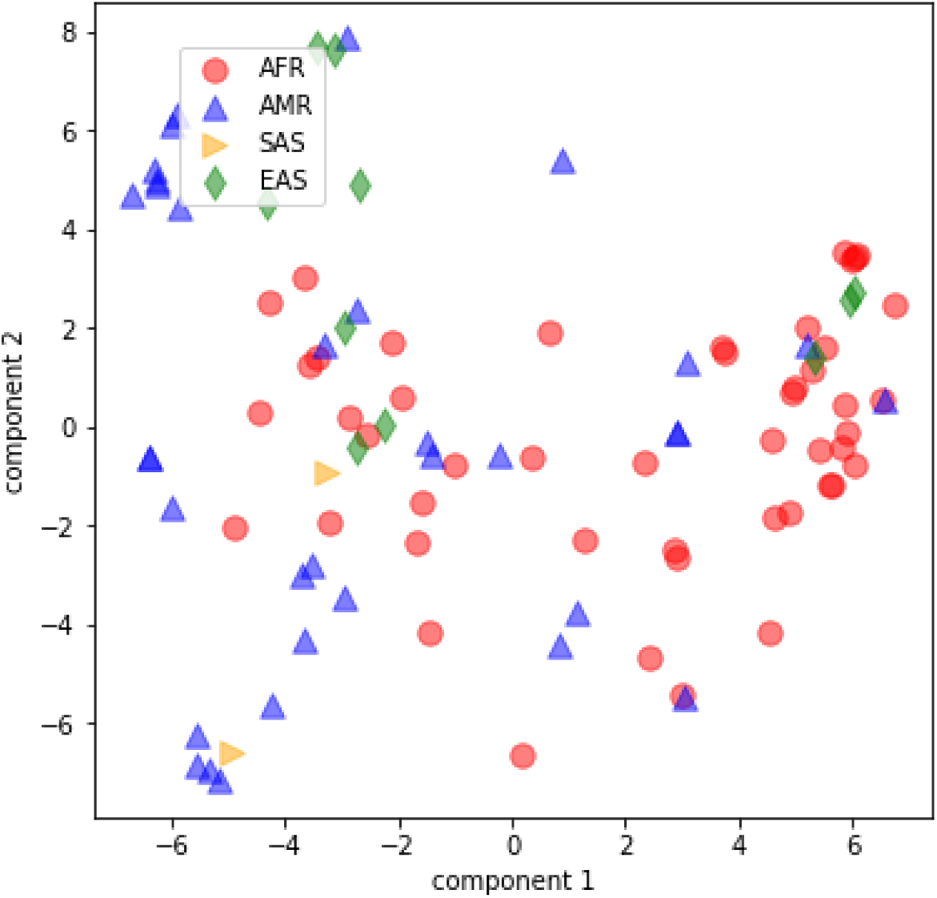
PCA plot for SNTG2 (Highest Entropy in the CMRG gene set)

**Supplementary Figure 6c:**
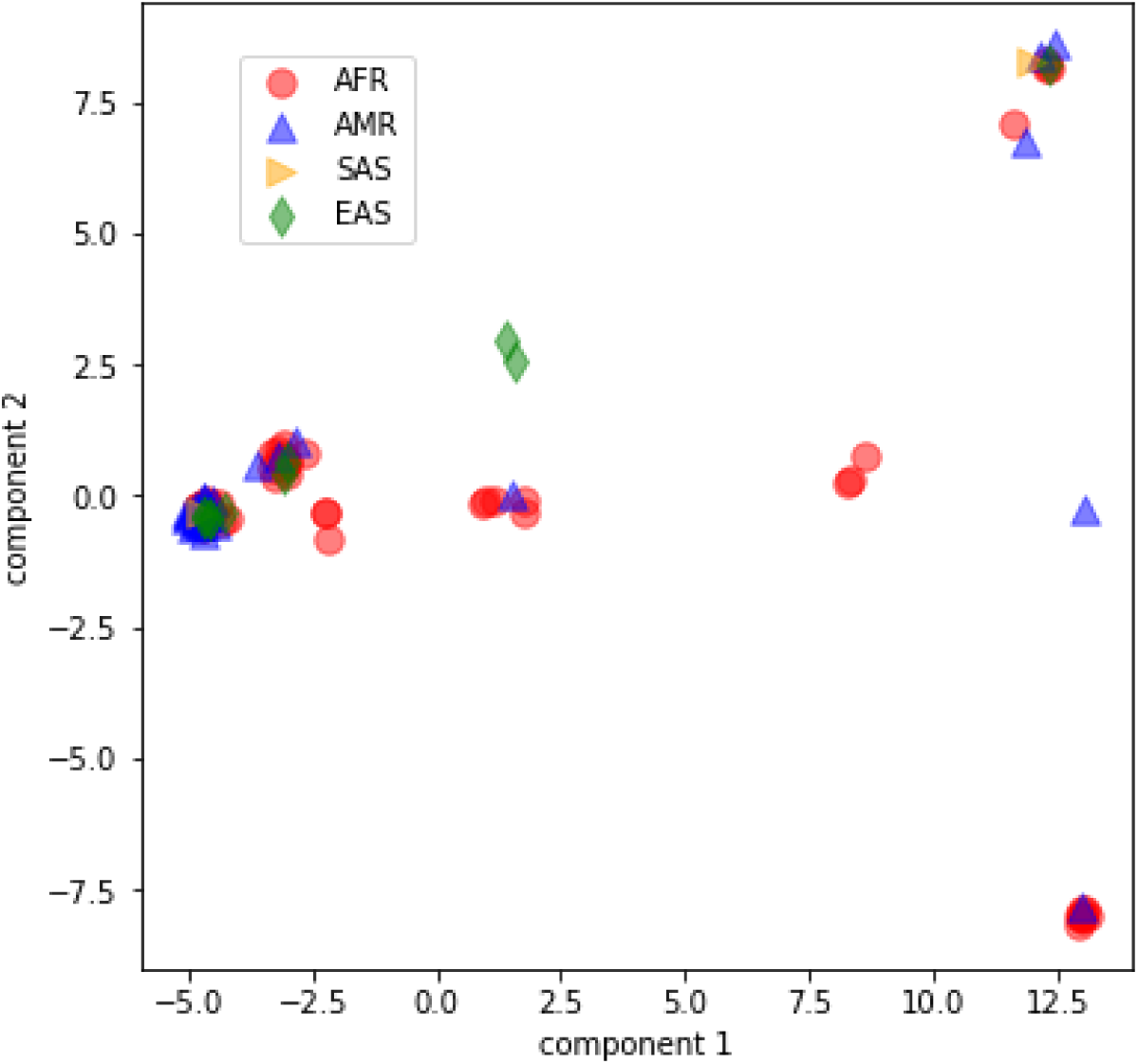
PCA plot for KMT2C (Highest Entropy in the CMRG gene set)

**Supplementary Figure 7.**
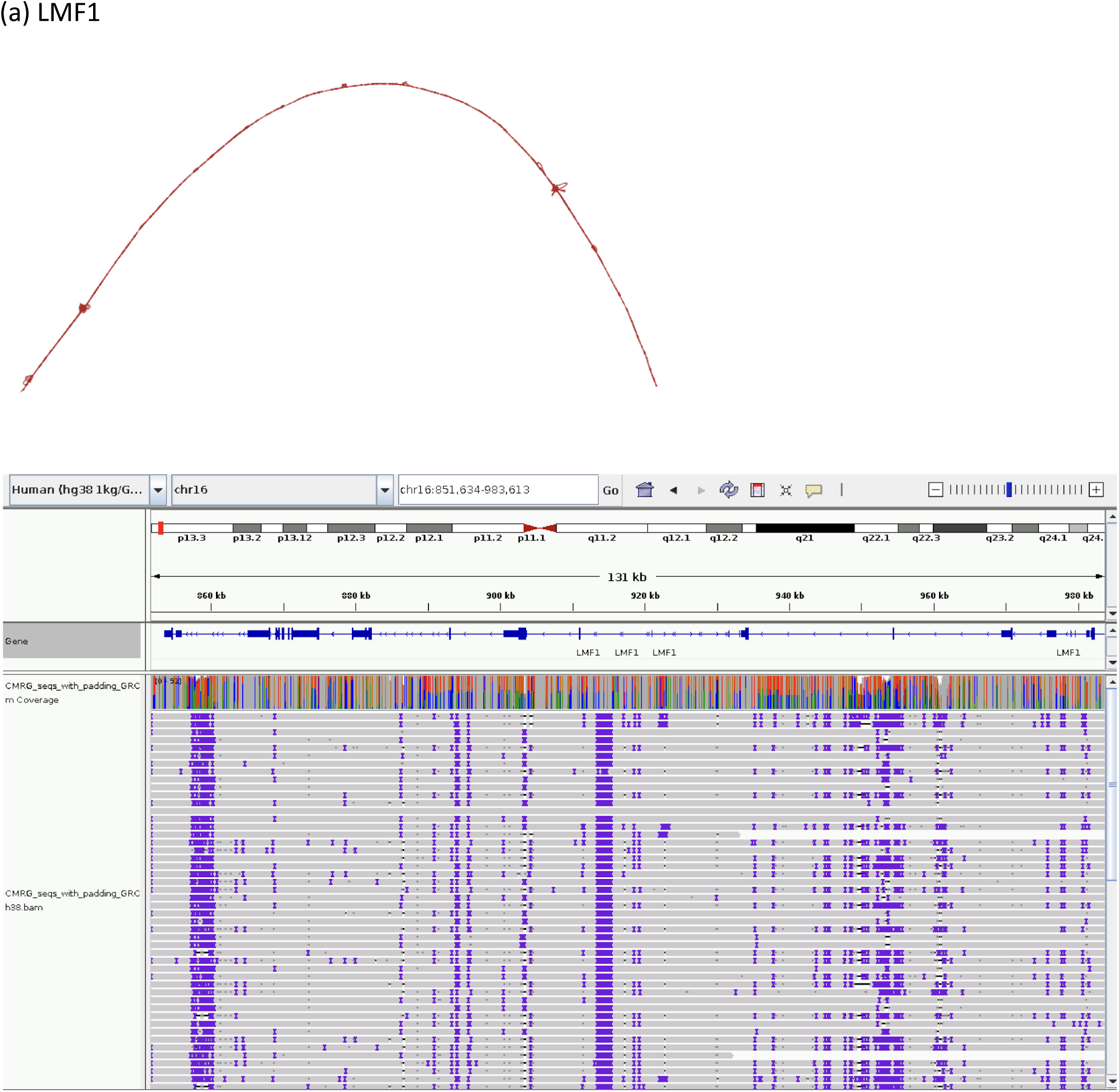
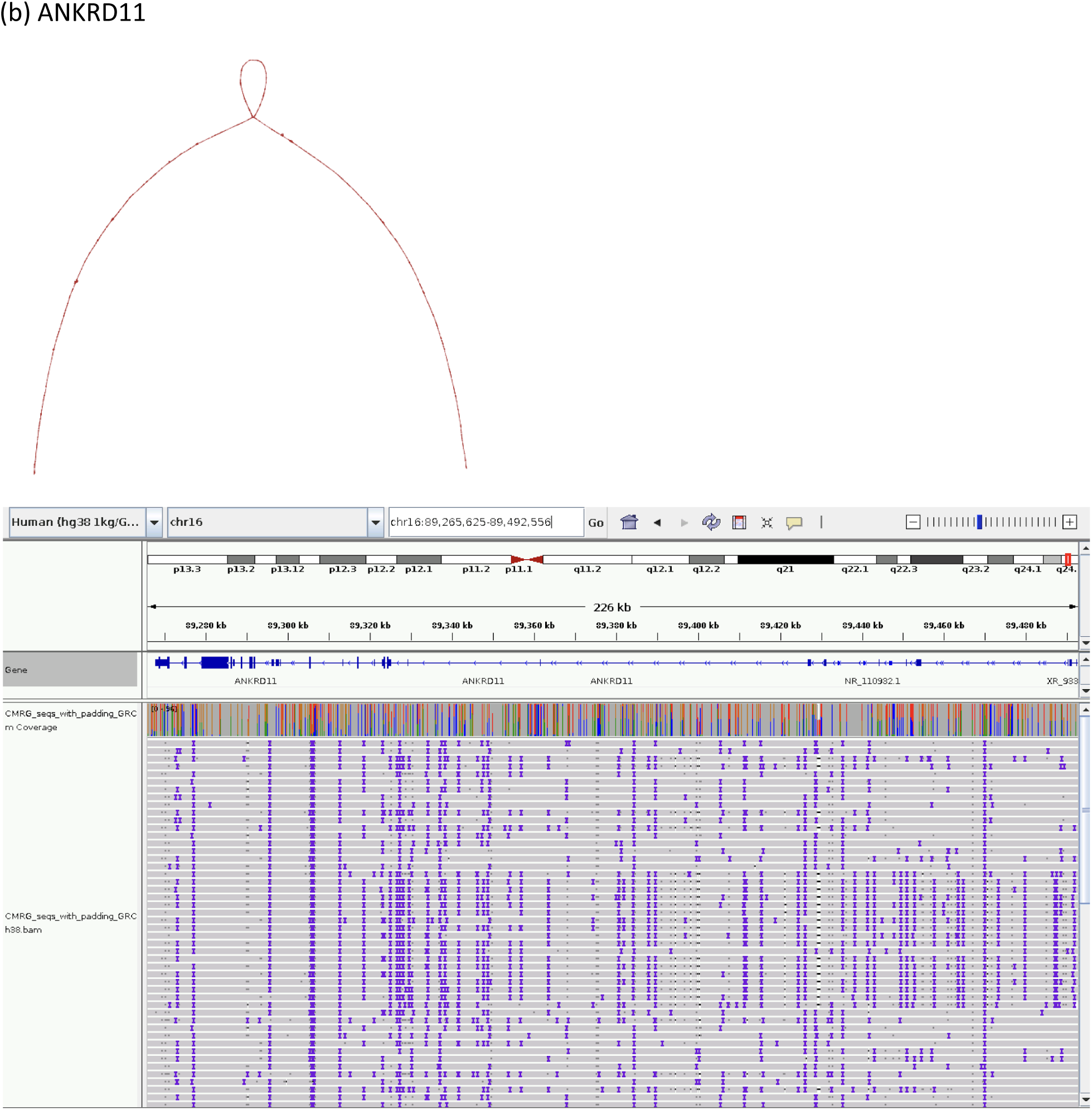
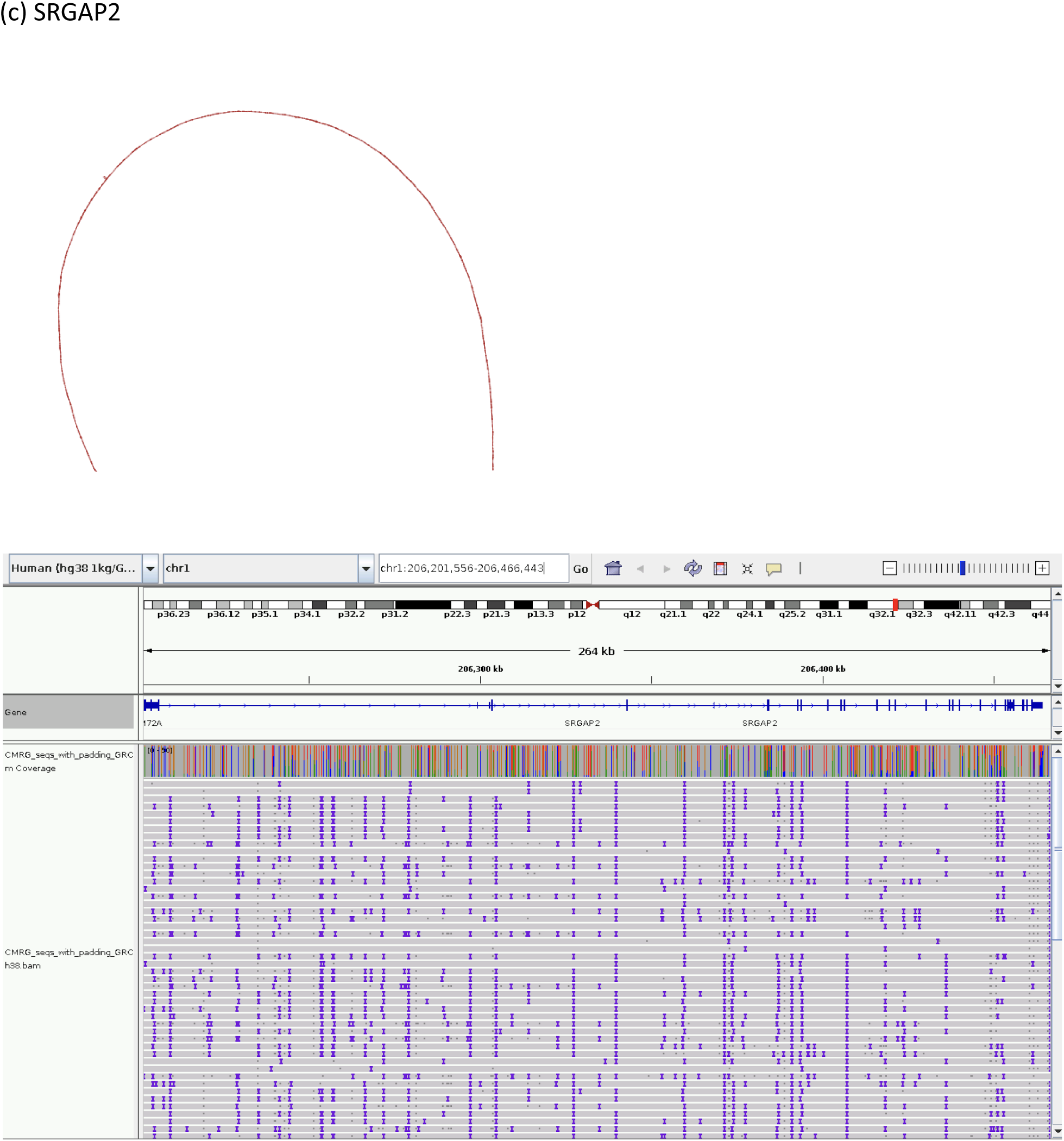
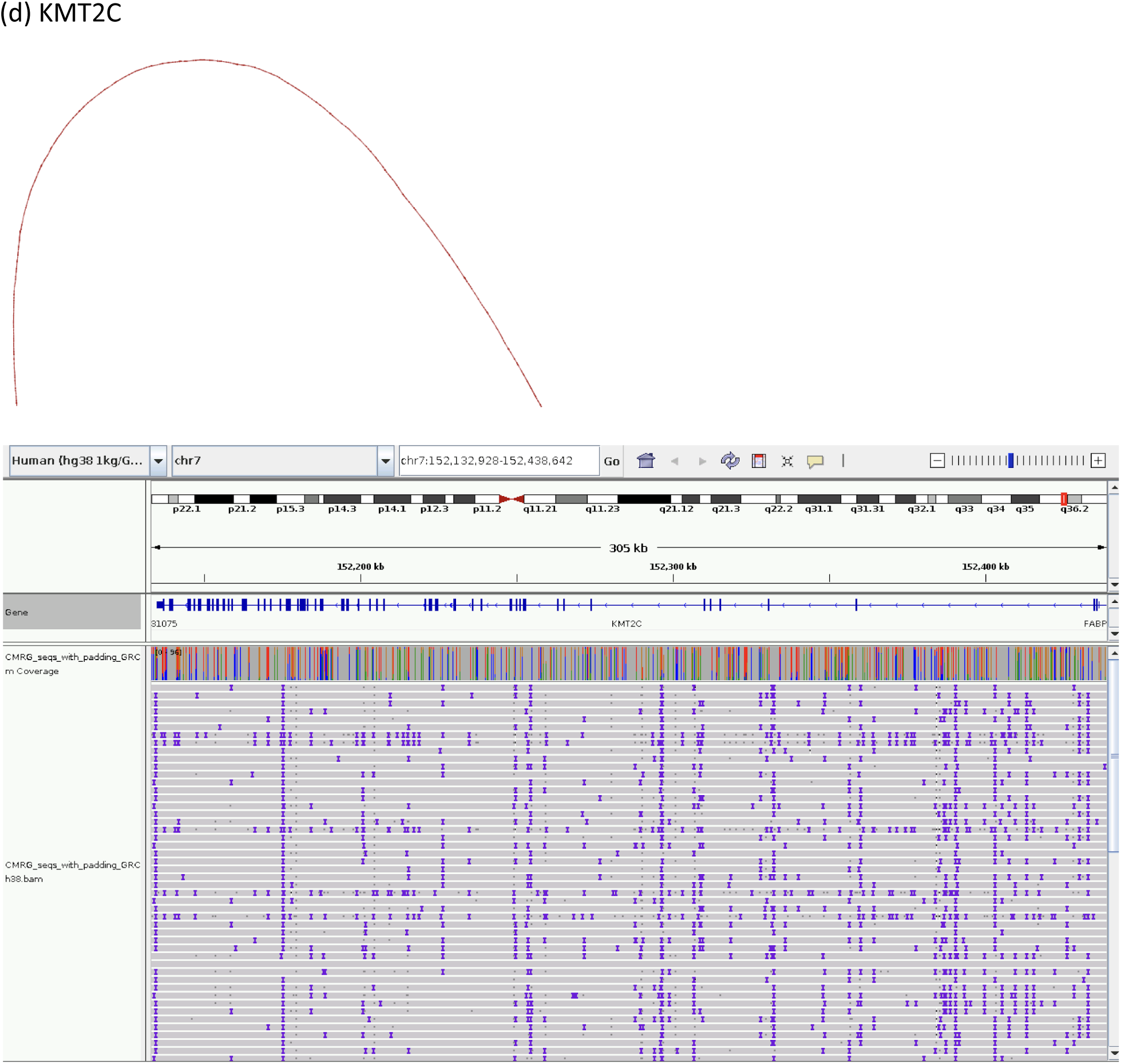
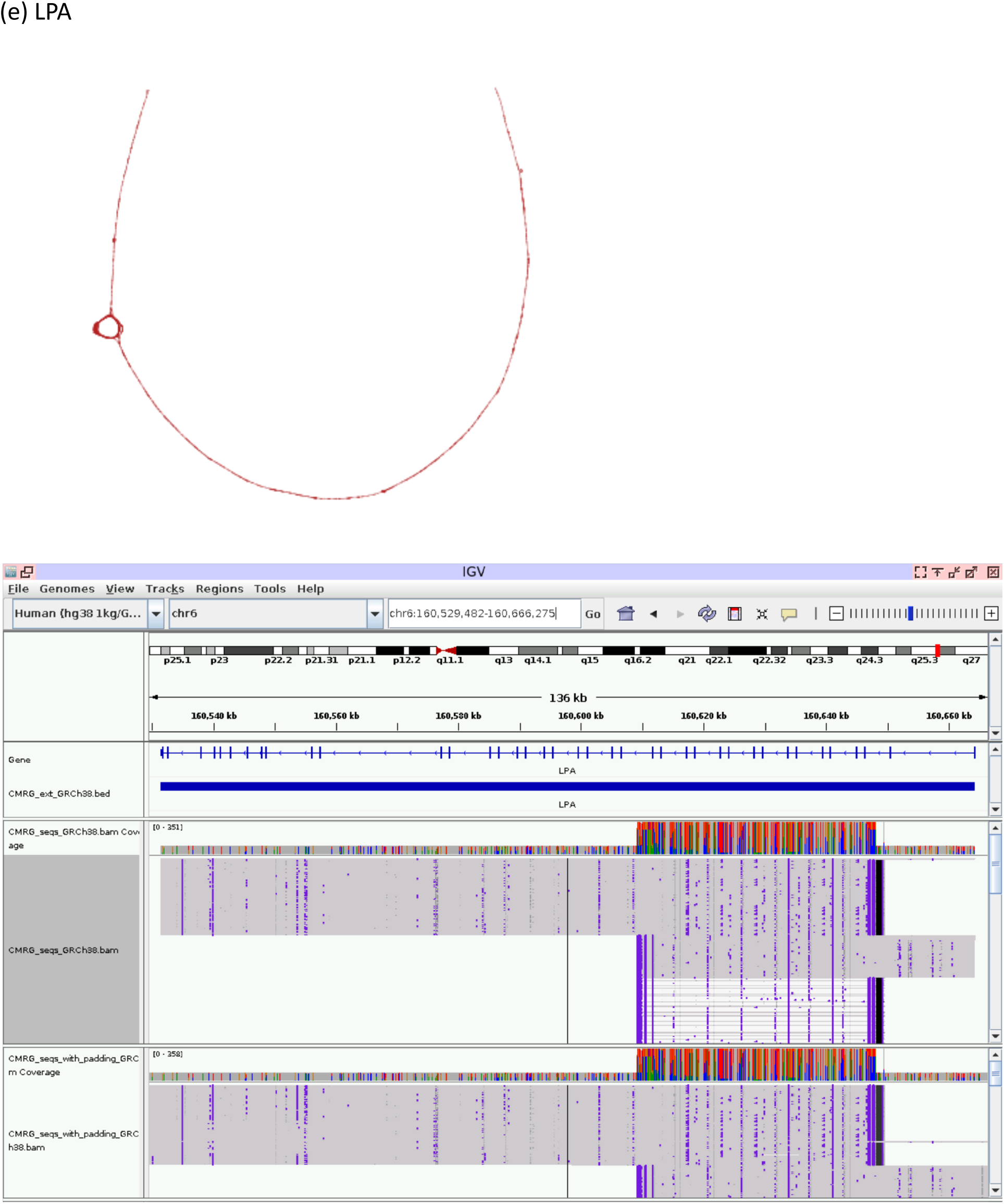
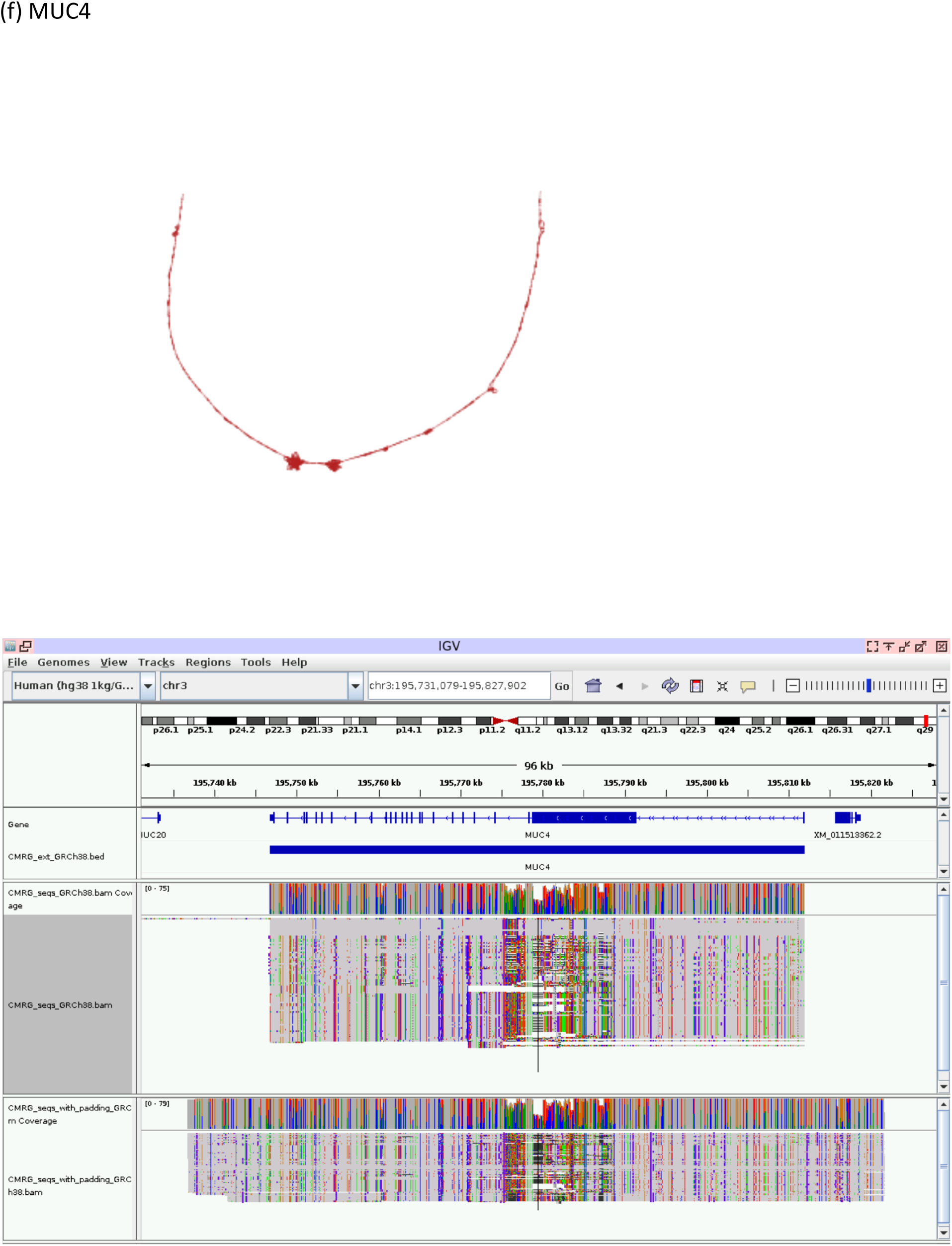
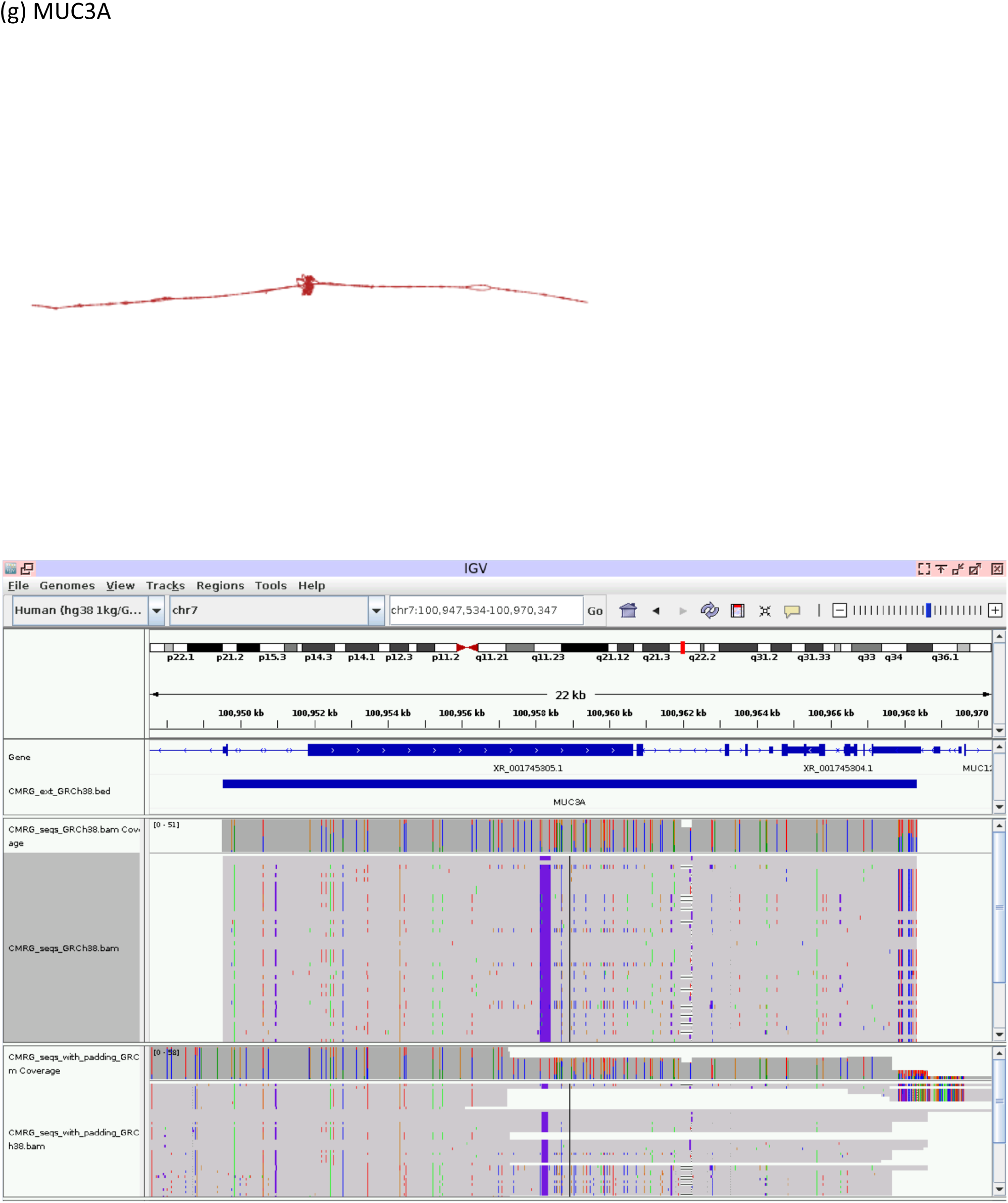
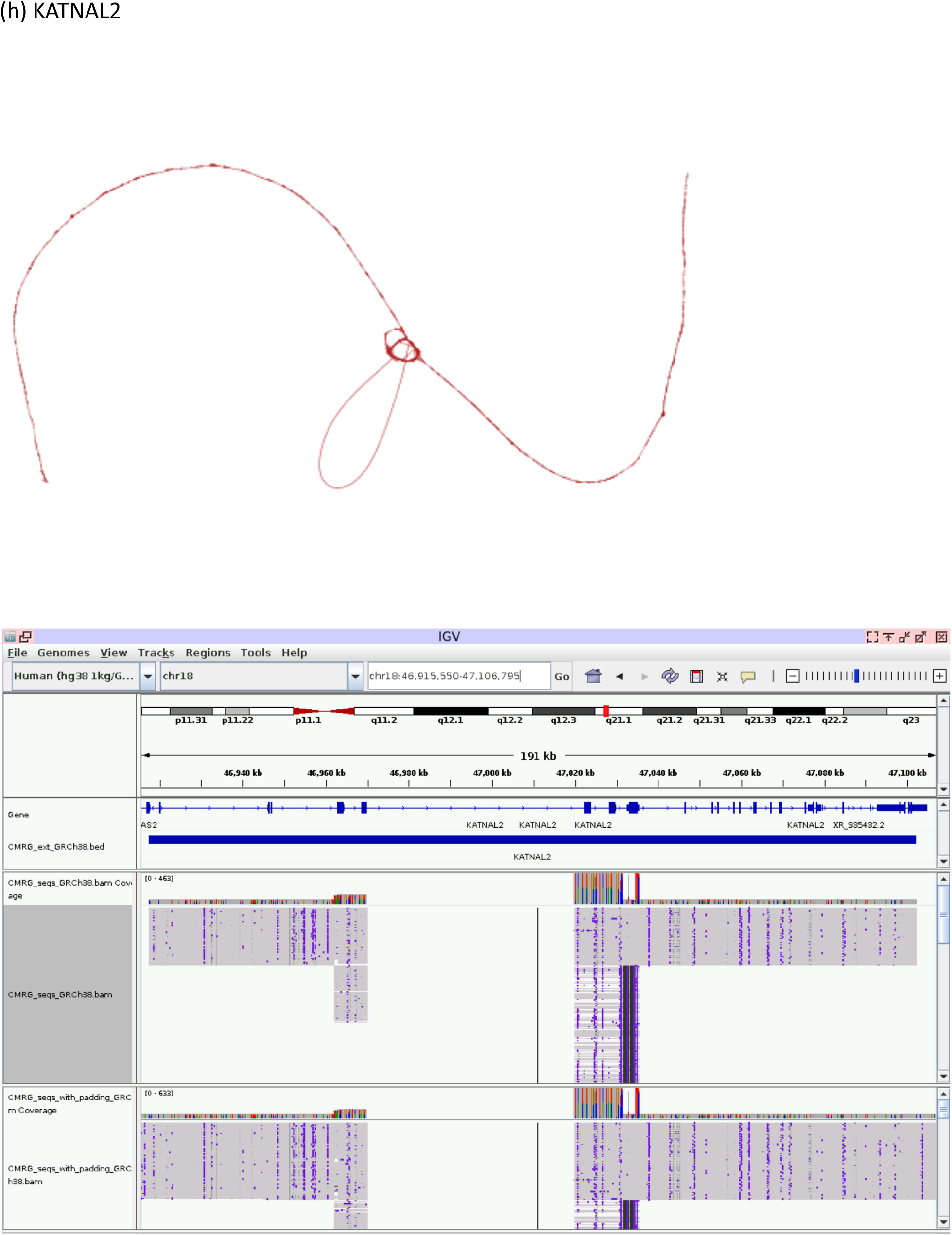
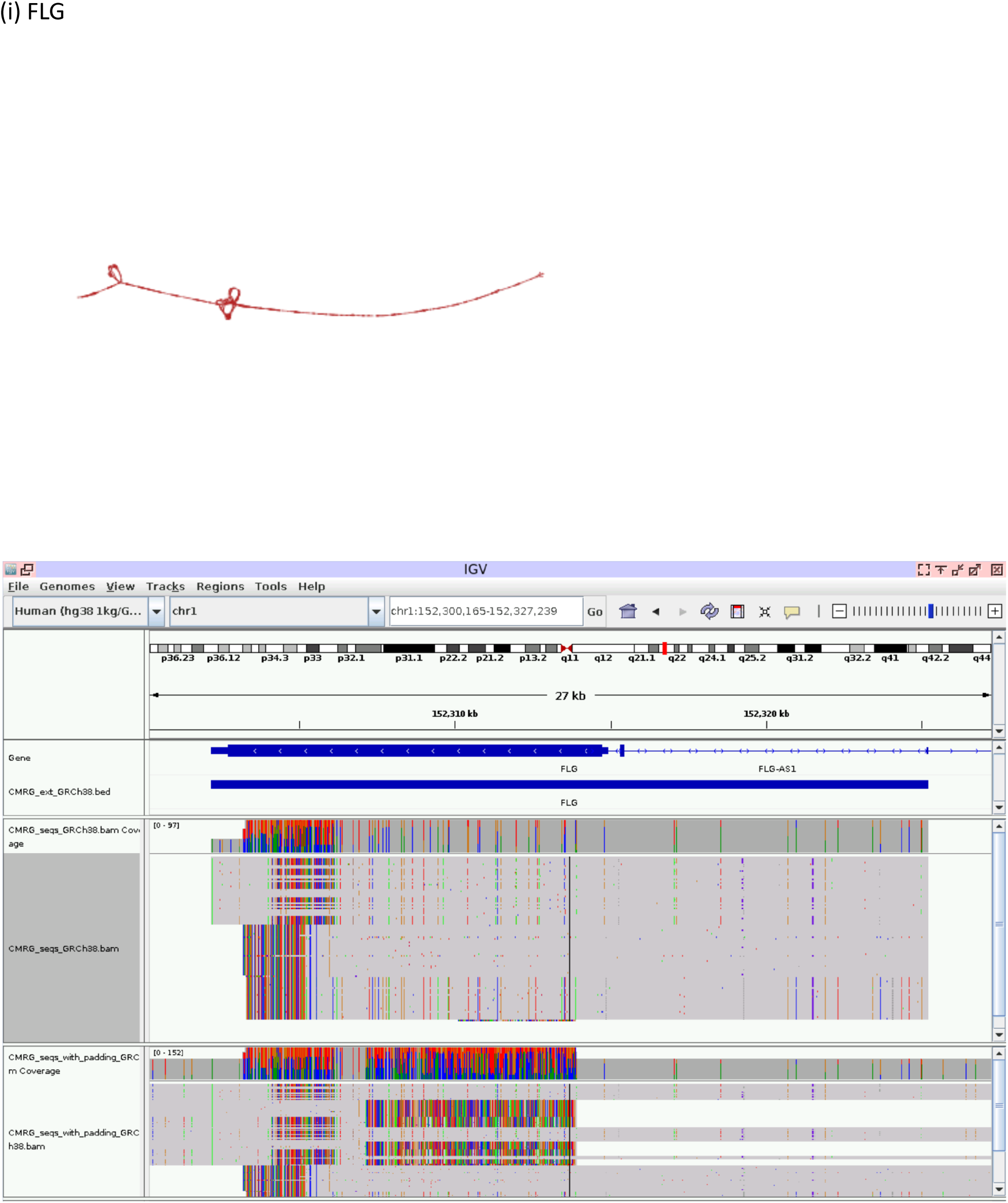
GIAB CMRG cases:

**Supplementary Figure 8.**
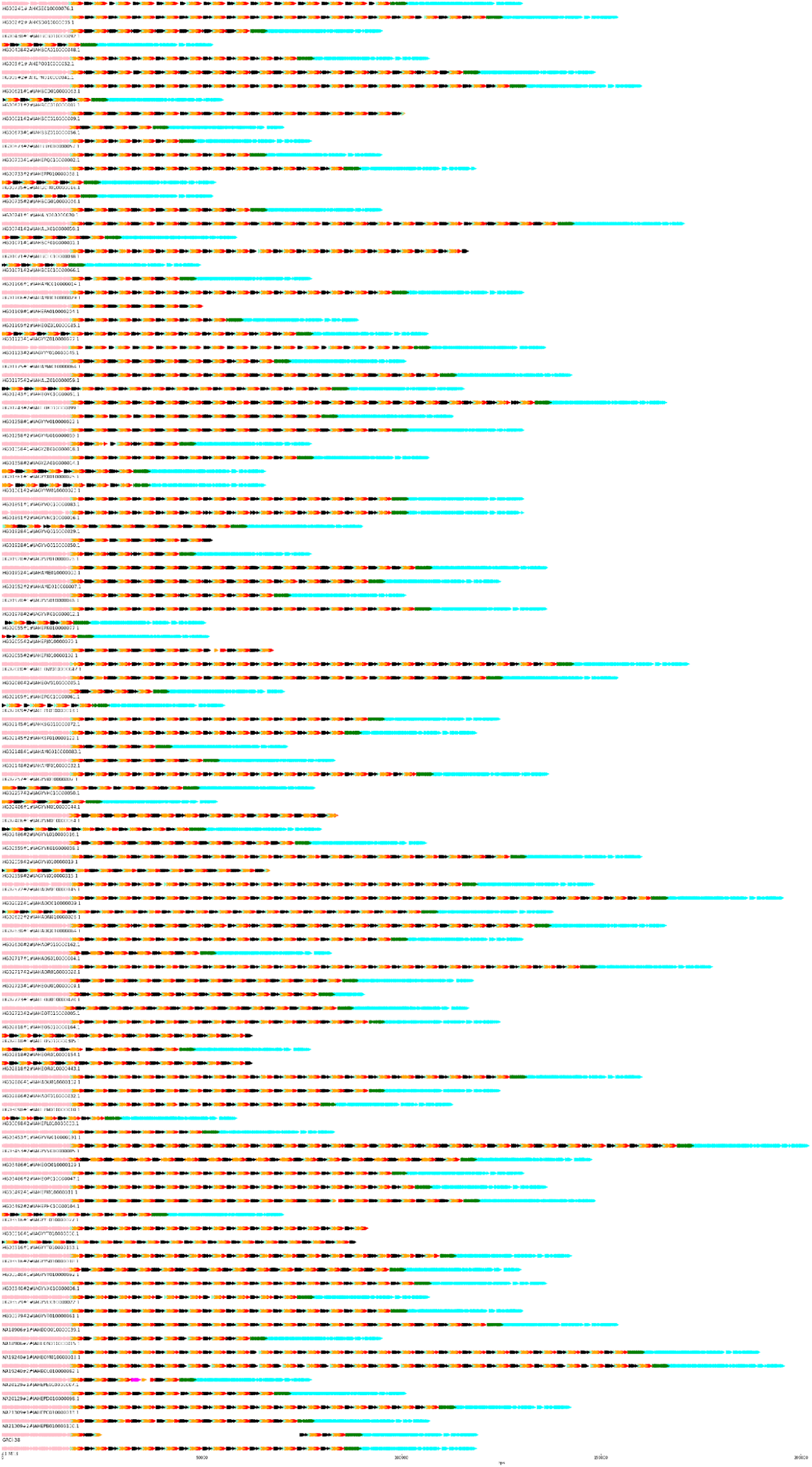
Principal bundle plot for KATNAL2: hg19 chr18: 44526786-44588614 showing different numbers of the repeat.

